# MOV10 inhibits SADS-CoV replication by enhancing TRIM24-mediated K63-linked TRAF3 ubiquitination and this inhibition is antagonized by viral N protein

**DOI:** 10.64898/2026.01.09.698592

**Authors:** Miaomiao Zeng, Dakai Liu, Jiyu Zhang, Liaoyuan Zhang, Hongyan Shi, Xin Zhang, Jianfei Chen, Xiuwen Li, Jialin Zhang, Tingshuai Feng, Xinwei Sun, Junyi Su, Zhaoyang Ji, Li Feng, Da Shi

**Author notes:** Corresponding author:* Li Feng, PhD, Da Shi, PhD. These authors contributed equally to this work (Miaomiao Zeng, Dakai Liu, Jiyu Zhang).

## Abstract

The global outbreak of SARS-CoV-2 has resulted in a renewed focus on coronaviruses with the potential for cross-species transmission and mutation risks. In particular, swine acute diarrhea syndrome coronavirus (SADS-CoV), potentially originating from intermediate horseshoe bat (*Rhinolophus affinis*), is a novel coronavirus that causes acute diarrhea, vomiting, and high mortality in suckling piglets. Innate immunity plays a crucial role in defending the host against invading pathogens. Previous studies have identified several host factors that inhibit SADS-CoV replication through innate immunity; however, research on SADS-CoV remains insufficient compared to that on other coronaviruses. In this study, the host factor Moloney leukemia virus 10 protein (MOV10) inhibited SADS-CoV replication by promoting interferon (IFN) production. Mechanistically, MOV10 exerted its antiviral effect primarily through its N-terminal domain, which regulated the TNF receptor-associated factor 3 (TRAF3)-mediated innate immune pathway. Specifically, we reveal that MOV10 enhanced tripartite motif-containing 24 (TRIM24)-mediated K63-linked ubiquitination of TRAF3, thereby promoting IFN production and inhibiting SADS-CoV replication. Furthermore, we identified the strategies by which SADS-CoV evaded innate immunity. The viral N protein inhibited the antiviral effect of MOV10 by disrupting interaction between MOV10 and TRAF3, and by promoting K48-linked polyubiquitination of TRAF3, leading to TRAF3 degradation via the ubiquitin-proteasome pathway. Collectively, our findings reveal the role of MOV10 in antiviral immunity and how SADS-CoV evades host immune defense.

## Introduction

Circulating coronaviruses in wildlife populations pose a substantial public health threat due to their zoonotic potential. Based on genetic and phenotypic characteristics, coronaviruses are classified into four distinct genera: *Alphacoronavirus*, *Betacoronavirus*, *Gammacoronavirus*, and *Deltacoronavirus*. In general, the former two infect only mammals, while the latter two mostly infect birds, but some of them can also infect mammals[1]. Several alphacoronaviruses, such as porcine epidemic diarrhea virus (PEDV), porcine transmissible gastroenteritis virus (TGEV), and SADS-CoV, cause significant gastroenteritis in neonatal piglets[2].

SADS-CoV is a novel coronavirus first identified in China in 2017[3]. The clinical signs of SADS-CoV infection in piglets include acute diarrhea, vomiting, and a mortality rate approaching 90%[4, 5]. Since its initial discovery, SADS-CoV has caused outbreaks in multiple provinces across China[4, 6], resulting in significant economic losses. SADS-CoV has recently been detected in swine populations in Vietnam, indicating that the virus has spread there from China[7]. This suggests that the virus has extended geographically, warranting global attention. The nucleocapsid (N) protein has the highest abundance in infected cells and is essential for viral transcription and assembly[8, 9]. In addition to viral function, the N protein is considered an important interferon (IFN) antagonist. For example, it performs this role during infection by severe acute respiratory syndrome coronavirus 2 (SARS-CoV-2)[10], severe acute respiratory syndrome coronavirus (SARS-CoV)[11], mouse hepatitis virus (MHV)[12], porcine deltacoronavirus (PDCoV)[13], and PEDV[14]. Although SADS-CoV has been shown to inhibit IFN-β production[15], the roles of its proteins in evading antiviral immunity and the underlying molecular mechanisms remain unclear.

Moloney leukemia virus 10 protein (MOV10), a 110-kDa RNA helicase, is a known component of cytoplasmic RNA processing centers called P-bodies[16]. Its helicase activity unwinds RNA in an ATP-dependent manner and in a 5’ to 3’ direction[17]. Additionally, MOV10 is an IFN-stimulated gene (ISG)[18, 19] that plays a crucial role in the innate immune response. During viral infection, viral RNA is recognized by host RNA sensors, such as retinoic acid inducible gene Ⅰ (RIG-Ⅰ), which triggers production of IFN via the MAVS-TRAF3-TBK1-IRF3 signaling pathway[20]. This induces MOV10 expression, thereby enabling it to target a wide range of RNA viruses through diverse mechanisms[21]. Ubiquitination, a key post-translational modification, plays a critical role in regulating the host IFN signaling pathway[22]. Upon RNA virus infection, the signaling molecules in this pathway, such as RIG-Ⅰ, mitochondrial antiviral signaling protein (MAVS; also known as IPS-1, VISA, and CARDIF), and TRAF3, undergo different types of ubiquitination by various E3 ubiquitin ligases and thus have different outcomes[23]. For example, K48-linked ubiquitination guides these molecules for proteasome degradation[23], whereas K63-linked ubiquitination promotes activation of downstream signaling and enhances transcription of IFN-α/β[24]. The balance between these opposing ubiquitination pathways generates a negative feedback loop to restrain IFN-I signaling. Consequently, activation of the IFN-I signaling pathway is finely tuned by dynamic regulation of the ubiquitination status of these signaling molecules by E3 ligases and deubiquitinating enzymes (DUBs).

In this study, we found that SADS-CoV infection induced expression of MOV10 *in vitro* and *in vivo*. MOV10 inhibited SADS-CoV replication mainly through its N-terminal domain. We also demonstrated that MOV10 enhanced the IFN-β response during SADS-CoV infection, with TRAF3 playing a critical role in the MOV10-regulated innate immune pathway. Additionally, MOV10 promoted K63-linked polyubiquitination of TRAF3 mediated by TRIM24. Furthermore, we investigated how SADS-CoV evaded the host immune system. The antiviral function of MOV10 was counteracted by viral N protein, which disrupted interaction of MOV10 with TRAF3, impairing the IFN signaling pathway. The N protein also directly targeted TRAF3 for K48-linked ubiquitination and degradation, inhibiting IFN production during the later stages of SADS-CoV infection. In summary, our findings highlighted the essential antiviral role of MOV10 in SADS-CoV infection and revealed how SADS-CoV manipulated the IFN signaling pathway to evade innate immune response. These insights provide a deeper understanding of the complex interaction between viral evasion strategies and host immune defenses.

## Results

### Induction of MOV10 expression by SADS-CoV infection *in vitro* and *in vivo*

The coronavirus N protein mediates the host innate immune response through multiple mechanisms, including modulation of host immune signaling pathways, interaction with host factors, and regulation of antiviral gene expression[31, 32]. To identify host factors that inhibit SADS-CoV immune evasion, we performed immunoprecipitation with Ab 3E9 against the SADS-CoV N protein, followed by LC-MS/MS analysis of the enriched samples (Fig. 1A). Among the identified host factors, MOV10 exhibited the highest fold change in protein expression (Fig. 1B). Co-IP assays confirmed this interaction (Fig. 1C and D). We then divided the N protein into three truncated regions to investigate the regions involved in the interaction (N1, N2, and N3) (Extended Data Fig. 1A). Our results demonstrated that MOV10 interacted with the N1 and N2 regions (Extended Data Fig. 1B and C). To investigate whether the interaction between the SADS-CoV N protein and MOV10 was dependent on RNase activity, we overexpressed both SADS-CoV N protein and MOV10 in HEK293T cells with or without RNase treatment. We observed a reduction in the interaction between SADS-CoV N protein and MOV10 in the presence of RNase A, suggesting that the interaction between SADS-CoV N and MOV10 was likely RNA-dependent (Fig. 1E). These results demonstrated that MOV10 may serve as a key host factor in mediating the host innate immune response to SADS-CoV N protein.

**Figure 1.**
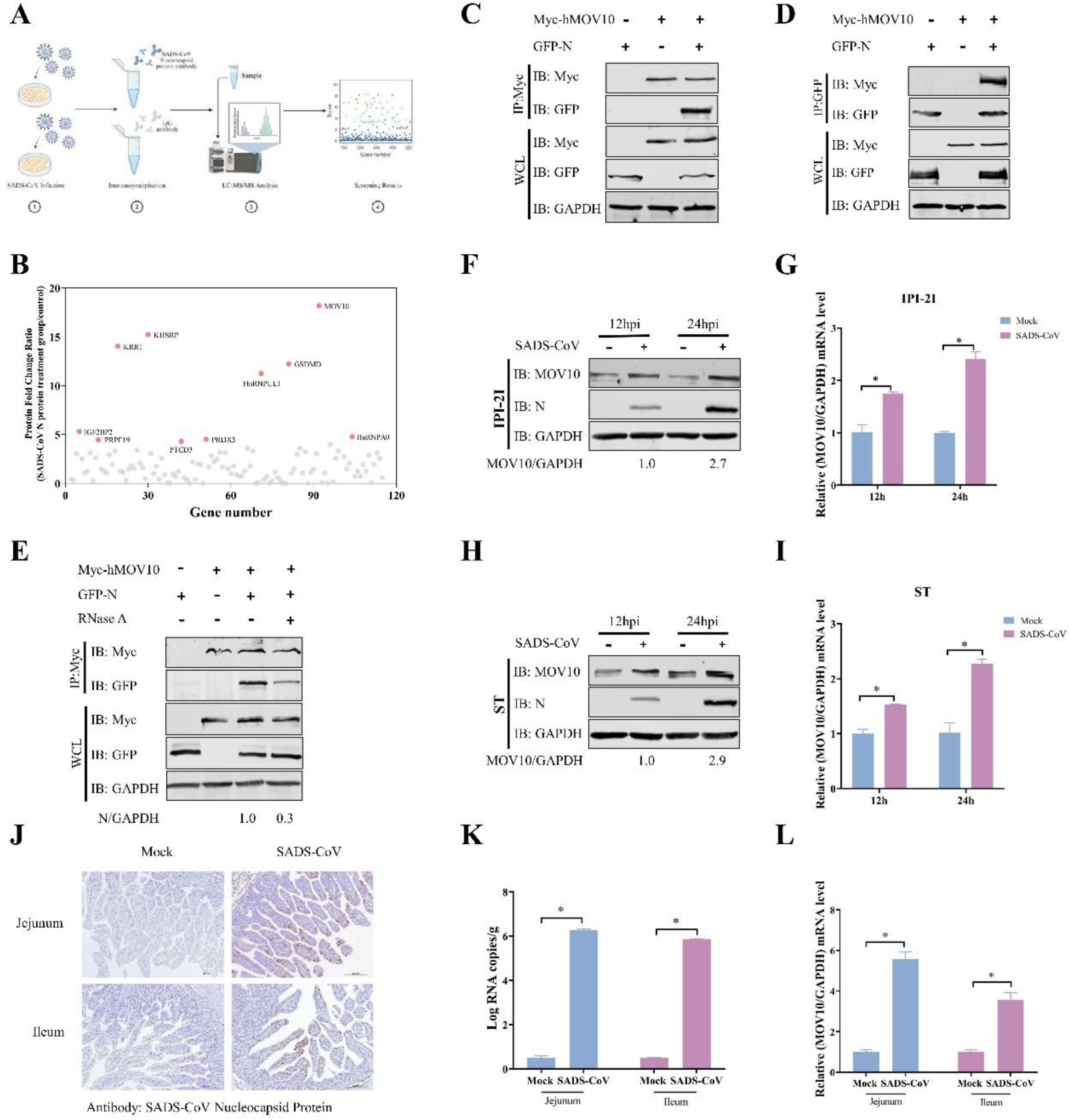
Induction of MOV10 expression by SADS-CoV infection *in vitro* and *in vivo*. (A) Flowchart illustrating the identification of host factors associated with SADS-CoV N protein. (B) Fold-change in protein expression (y axis) and gene number (x axis) of host factors identified in Vero E6 cells infected with SADS-CoV. (C-D) MOV10 interacted with the SADS-CoV N protein. HEK293T cells were transfected or co-transfected with plasmids encoding Myc-hMOV10 and GFP-N. Cell lysates were subjected to immunoprecipitation using Myc (C) or GFP (D) Abs. (E) Interaction between SADS-CoV N and MOV10 was likely RNA-dependent. Plasmids encoding Myc-hMOV10 and GFP-N were transfected or co-transfected into HEK293T cells in the presence or absence of RNase A (100 µg/mL). Cell lysates were subjected to immunoprecipitation using Myc Abs. The density of SADS-CoV N protein bands relative to GAPDH was calculated, with the values of each negative control group normalized to 1. (F-I) Expression of MOV10 was upregulated following SADS-CoV infection. IPI-2I and ST cells were infected with SADS-CoV (MOI = 0.1) and harvested at 12 and 24 hpi. Cell lysates were analyzed by western blotting (F and H) using Abs against MOV10 and the SADS-CoV N protein. GAPDH was used as a protein loading control. The density of MOV10 bands relative to GAPDH was calculated, with the values of each negative control group normalized to 1. MOV10 mRNA levels were quantified by qRT-PCR (G and I). (J) Representative photomicrographs of viral antigen immunochemical staining in SADS-CoV-infected and uninfected jejunum and ileum tissues (Bar: 200 µm). The virus in intestinal tissue cells was immunostained using mAb 3E9 against SADS-CoV N protein. (K) Viral RNA copy numbers in the jejunum and ileum samples were analyzed by qRT-PCR. (L) MOV10 levels in the jejunum and ileum of SADS-CoV-infected and uninfected piglets at 36 hpi were quantified by qRT-PCR. The mean and SD of the results from three independent experiments are shown (\**P* < 0.05).

To confirm the function of MOV10 in the replication of SADS-CoV, we first assessed expression of MOV10 *in vitro* and *in vivo* following SADS-CoV infection. IPI-2I and ST cells were infected with SADS-CoV (MOI=0.1) for 12 and 24 h. We observed that both the protein (Fig. 1F and H) and mRNA (Fig. 1G and I) levels of MOV10 in SADS-CoV-infected cells showed an obvious increase compared to the control group. Three-day-old SPF piglets were orally challenged with SADS-CoV, and unchallenged piglets served as mock controls. Jejunal and ileal tissues from both groups were harvested at 36 hpi, and successful infection was confirmed by IHC and qRT-PCR (Fig. 1J and K). Notably, the MOV10 mRNA levels were also significantly higher in the jejunum and ileum of SADS-CoV-infected piglets compared to controls (Fig. 1L). These findings revealed that SADS-CoV infection upregulated MOV10 expression *in vitro* and *in vivo*.

### MOV10 is a host restriction factor for SADS-CoV replication

To further elucidate the role of MOV10 in SADS-CoV replication, IPI-2I cells overexpressing porcine MOV10 were infected with SADS-CoV. We found that the levels of viral N protein in SADS-CoV-infected IPI-2I cells were significantly reduced (Fig. 2A and B). Additionally, a marked decrease in viral titer was also observed (Fig. 2C). We next extended our findings to ST cells. It is not surprising that SADS-CoV N protein expression (Fig. 2D), mRNA levels (Fig. 2E), and viral titer (Fig. 2F) showed an apparently decrease. Our findings demonstrated that overexpression of MOV10 suppressed the replication of SADS-CoV in a dose-dependent manner. Furthermore, three siRNAs specifically designed to target porcine MOV10 were employed to downregulate its expression in IPI-2I cells. Among these, siMOV10-1 and siMOV10-3 resulted in the most significant downregulation of MOV10 levels (Extended Data Fig. 2A and B). We subsequently found that knockdown of MOV10 resulted in a pronounced increase in viral N protein and mRNA levels (Fig. 2G and H) and enhanced the release of infectious virus (Fig. 2I). To further confirm MOV10 as a host restriction factor, we generated MOV10-deficient IPI-2I cells (MOV10^−/−^) through CRISPR-mediated genome editing (Extended Data Fig. 3A-C). As shown in Fig. 2J-L, we observed significant upregulation of viral N protein expression and viral titers in MOV10^−/−^ cells compared with wild type (WT) cells. Collectively, these results indicated that MOV10 is a host restriction factor for SADS-CoV replication.

**Figure 2.**
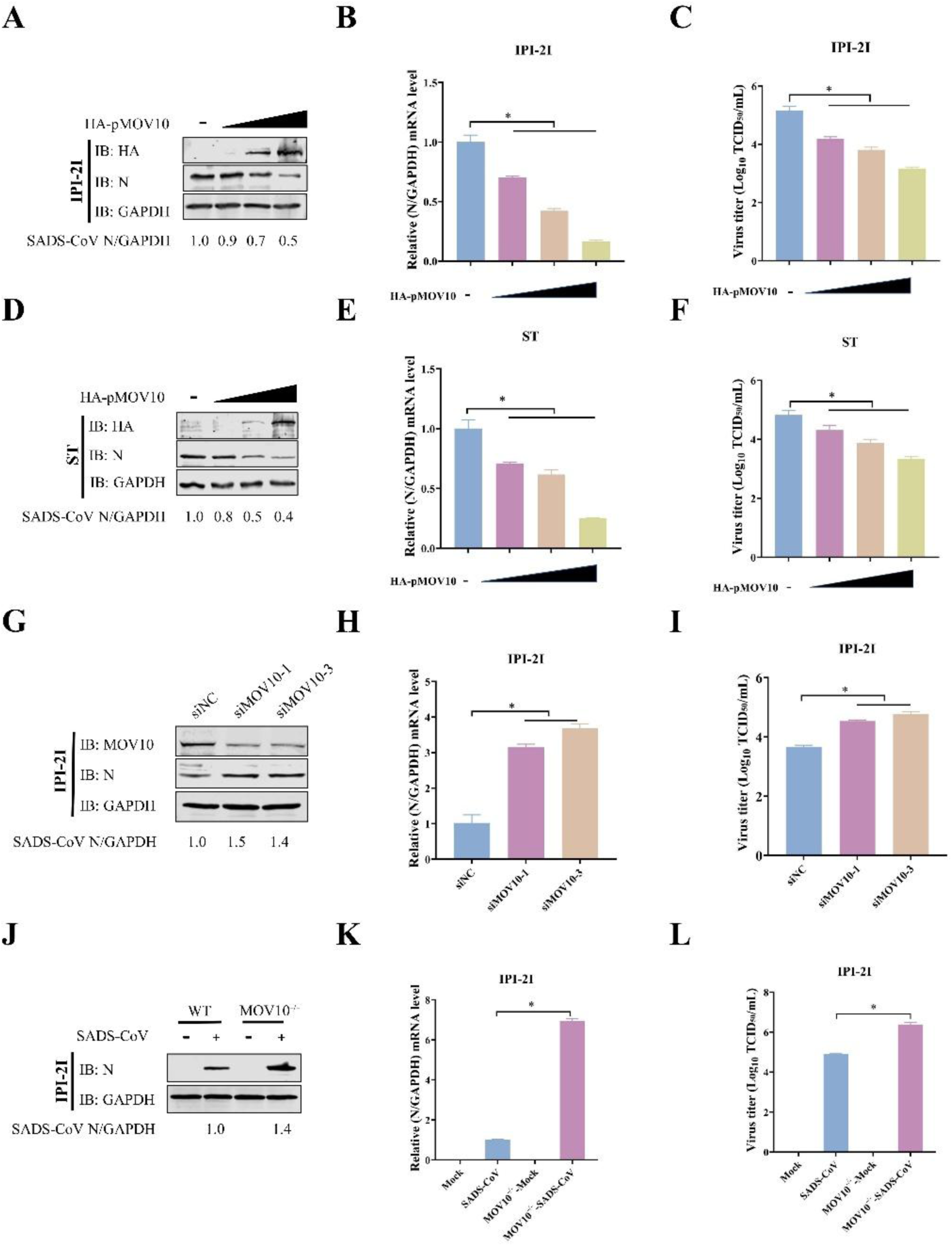
MOV10 is a host restriction factor for SADS-CoV infection. (A-F) Overexpression of MOV10 inhibited SADS-CoV replication. IPI-2I and ST cells were transfected with varying concentrations (0, 1, 3, 5 μg) of full-length porcine MOV10 (HA-pMOV10) for 24 h, followed by infection with SADS-CoV (MOI = 0.1) for an additional 24 h. Cell lysates were analyzed by western blotting (A and D) using Abs against the HA tag, SADS-CoV N protein and GAPDH. The density of the N protein band relative to GAPDH was calculated, with values from each negative control group normalized to 1. mRNA levels of SADS-CoV N protein were quantified by qRT-PCR (B and E). Viral titers were determined using TCID_50_ assays (C and F). (G-I) Knockdown of MOV10 enhanced replication of SADS-CoV. IPI-2I cells were transfected with siMOV10-1, siMOV10-3, or negative control (siNC) at 50 nM for 48 h and then infected with SADS-CoV (MOI = 0.1) for 24 h. Lysates were analyzed by western blotting (G) using Abs against MOV10, SADS-CoV N protein and GAPDH. Band density for SADS-CoV N/GAPDH was calculated, with values from the siNC group standardized to 1. mRNA levels of SADS-CoV N detected by qRT-PCR (H). Viral titers were determined using TCID_50_ assays (I). (J-L) Knockout of MOV10 enhanced SADS-CoV replication. WT and MOV10^−/−^ IPI-2I cells were infected with SADS-CoV (MOI = 0.1) or uninfected for 24 h. Lysates were analyzed by western blotting (J). Density of SADS-CoV N bands relative to GAPDH was calculated using grayscale analysis. mRNA levels of SADS-CoV N protein were detected by qRT-PCR (K). Viral titers were determined using TCID_50_ assays (L). Mean and SD of the results from three independent experiments are shown (\**P* < 0.05).

### MOV10 inhibits SADS-CoV replication mainly through its N-terminal domain

MOV10 is an RNA helicase that contains an N-terminal CH-rich domain and a C-terminal helicase domain (Fig. 3A). The C-terminus of MOV10 (524-912aa) contains seven helicase motifs (I, Ia, II, III, IV, V and VI) required for its helicase activity, which have a DEAG fingerprint structure. Therefore, it is classified into the SF-1 family and its N-terminus (93-305 aa) contains a Cys-His-rich (CH) domain (Fig. 3B)[33, 34]. Wang et al. demonstrated that MOV10 inhibits Middle East respiratory syndrome coronavirus (MERS-CoV) through its helicase activity[35]. To determine whether MOV10-suppresses SADS-CoV replication by a similar mechanism, we infected IPI-2I cells transfected with either an N-terminal or C-terminal plasmid with the virus. As shown in Fig. 3C and D, full-length MOV10 and its N-terminal domain significantly reduced the levels of N protein and mRNA, whereas the C domain only resulted in a minor reduction. These findings were further confirmed by TCID_50_ assays (Fig. 3E). Moreover, similar results were observed in ST cells, supporting the consistency of our observations (Fig. 3F-H). Interestingly, these results reveled that MOV10 primarily inhibits SADS-CoV replication through its N-terminal domain, rather than relying on its helicase activity. This finding suggested that, despite both viruses belonging to the same family of coronaviruses, the mechanism by which MOV10 inhibits SADS-CoV differs significantly from that of MERS-CoV.

**Figure 3.**
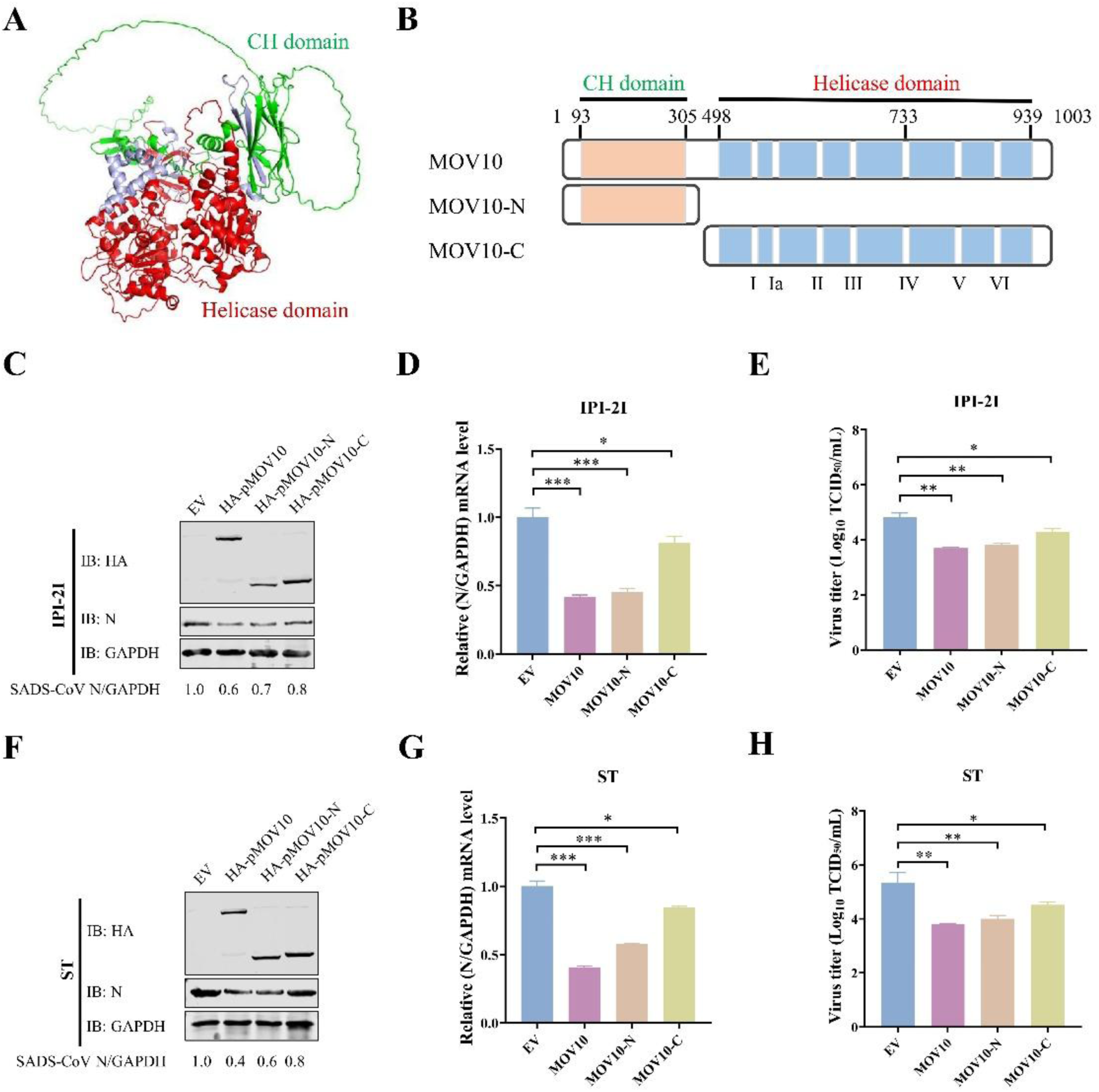
MOV10 suppressed SADS-CoV replication independent of its RNA helicase activity. (A) Structural of CH (green) and helicase (red) domains of porcine MOV10. (B) Schematic diagram of truncated MOV10 mutants. Numbers indicate amino acid positions. The orange boxes represent CH domain. The blue boxes to the left and right represent helicase domains, respectively. The positions of the seven helicase motifs are indicated under the third bar. (C-H) The N-terminus of MOV10 was required for inhibition of SADS-CoV replication. IPI-2I and ST cells were transfected with HA-pMOV10 or its truncations (HA-pMOV10-N and HA-pMOV10-C) for 24 h, followed by infection with SADS-CoV (MOI = 0.1) for an additional 24 h. Cell lysates were analyzed by western blotting (C and F). The density of SADS-CoV N bands relative to GAPDH was calculated using grayscale analysis. mRNA levels of SADS-CoV N was assessed by qRT-PCR (D and G). Viral titers were determined using TCID_50_ assays (E and H). The mean and SD of the results from three independent experiments are shown (\**P* < 0.05; \*\**P* < 0.01; \*\*\**P* < 0.001).

### MOV10 positively regulates lFN-β response upon SADS-CoV infection

Previous studies have indicated that MOV10 functions as an ISG^[18, 36]^, but its role as an ISG in porcine intestinal cells, such as IPI-2I cells, remains unclear. We subsequently assessed expression of MOV10 and ISG15 in IPI-2I and HEK293T cells in response to stimulation with 100 and 1000 ng/µL IFN-β. As expected, the levels of MOV10 and ISG15 were dramatically elevated in both cell lines treated with IFN-β compared to the control cells (Fig. 4A). In addition, both MOV10 and ISG15 reached their peak levels approximately 6∼8 h after treatment with 100 ng/µL IFN-β (Fig. 4B). These results shown that MOV10 is an ISG in IPI-2I and HEK293T cells. To further investigate the role of MOV10 in innate immunity, we analyzed its function by stimulation with RIG-I receptor ligands, such as SeV or poly(I:C)[37, 38]. We first investigated the effect of MOV10 on ISRE or IFN-β expression following SeV infection. Results revealed that MOV10 markedly enhanced SeV-triggered activation of both the ISRE and IFN-β (Fig. 4C and D). We then found that MOV10 enhanced the levels of IFN genes, including IFN-β and ISG15, in HEK293T cells triggered by SeV and poly(I:C) (Fig. 4E). Additionally, MOV10 also upregulated them in SADS-CoV infected IPI-2I and HEK293T cells (Fig. 4F).

**Figure 4.**
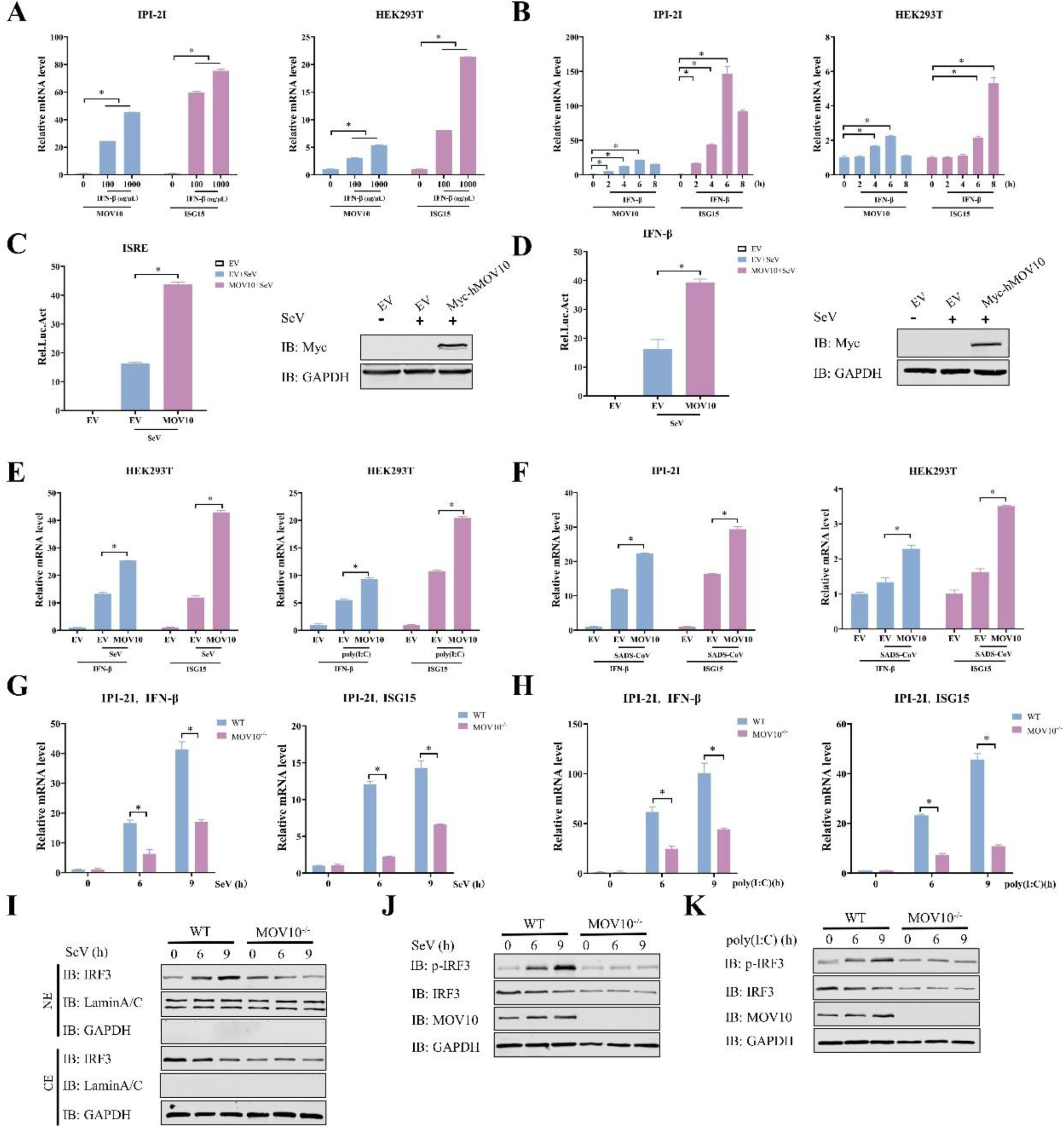
MOV10 positively regulated IFN-β response. (A and B) MOV10 was an ISG in IPI-2I and HEK293T cells. IPI-2I and HEK293T cells were treated with porcine or human recombinant IFN-β (0, 100, and 1000 ng/µL) for 8 h. mRNA levels of ISG15 and MOV10 were then measured by qRT-PCR (A). IPI-2I and HEK293T cells were treated with recombinant IFN-β (100 ng/µL) for 0, 2, 4, 6, and 8 h, and mRNA levels of MOV10 and ISG15 were measured by qRT-PCR (B). (C and D) HEK293T cells were transfected with Myc-hMOV10 or empty vector (EV) for 24 h, followed by infection with SeV for 9 h. ISRE (C) or IFN-β (D) were detected by luciferase reporter assay. Expression of MOV10 and GAPDH was determined by western blotting. (E) HEK293T cells were transfected with Myc-hMOV10 or EV for 24 h and then infected with SeV or stimulated with 10 µg/mL poly(I:C) for 9 h. mRNA levels of IFN-β and ISG15 were assessed by qRT-PCR. (F) IPI-2I and HEK293T cells were transfected with Myc-hMOV10 or EV for 24 h and infected with SADS-CoV (MOI = 0.1) for an additional 24 h. mRNA levels of IFN-β and ISG15 were analyzed by qRT-PCR. (G and H) WT and MOV10^−/−^ IPI-2I cells were infected with SeV or stimulated with 10 µg/mL poly(I:C) for 6 and 9 h. mRNA levels of IFN-β and ISG15 were quantified by qRT-PCR. (I) WT and MOV10^−/−^ IPI-2I cells infected with SeV were harvested at 0, 6 and 9 h. Protein expression of IRF3, LaminA/C, and GAPDH in cytoplasmic (CE) and nuclear (NE) fractions was analyzed by western blotting. (J and K) WT and MOV10^−/−^ IPI-2I cells were infected with SeV or stimulated with 10 µg/mL poly(I:C) for 0, 6 and 9 h. Western blotting was performed using Abs against p-IRF3, IRF3, MOV10, and GAPDH. Mean and SD of the results from three independent experiments are shown (\**P* < 0.05).

Given that MOV10 positively regulated the IFN response, it was imperative to investigate whether knockout of MOV10 also affected this response. As shown in Figure 4G and H, WT IPI-2I cells exhibited higher activation of IFN-β and ISG15 than MOV10^−/−^ cells in response to SeV infection or poly(I:C) stimulation. The data suggest that MOV10 induced activation of the IFN-I pathway. To further validate these findings, we assessed the effect of MOV10 on phosphorylation of IRF3, a key marker of IFN-I pathway activation, following SeV infection or poly(I:C) treatment. We found that SeV-induced IRF3 nuclear translocation was dramatically suppressed in MOV10^−/−^ cells compared to WT cells (Fig. 4I). Consistently, MOV10 deficiency markedly impaired phosphorylation of IRF3 in response to SeV infection or poly(I:C) stimulation (Fig. 4J and K). Thus, these results demonstrate that MOV10 is important for host defense against viral infection by mediating IFN-β signaling activation.

### TRAF3 plays a crucial role in the MOV10-regulated innate immune pathway

RIG-I-mediated innate immune signaling is critical for host defense against RNA virus infection [39, 40]. However, previous studies have reported that MOV10 against RNA virus infection by promoting the IFN-I response through an IKKε-mediated RNA sensing pathway, independent of the RIG-I–MAVS pathway[41], we were prompted to investigate its potential role against SADS-CoV by assessing its interaction with upstream regulators of IKKε, including RIG-I, MDA5, MAVS, and TRAF3. Notably, MOV10 was clearly associated with TRAF3, while no interaction was detected with RIG-I, MDA5 or MAVS (Fig. 5A). Interaction between MOV10 and TRAF3 was further predicted by AlphaFold3 (Fig. 5B). This interaction was confirmed by both confocal microscopy and Co-IP assays (Fig. 5C-E). These findings suggest that MOV10 modulated TRAF3-dependent signaling in the innate immune pathway, further influencing downstream immune responses.

**Figure 5.**
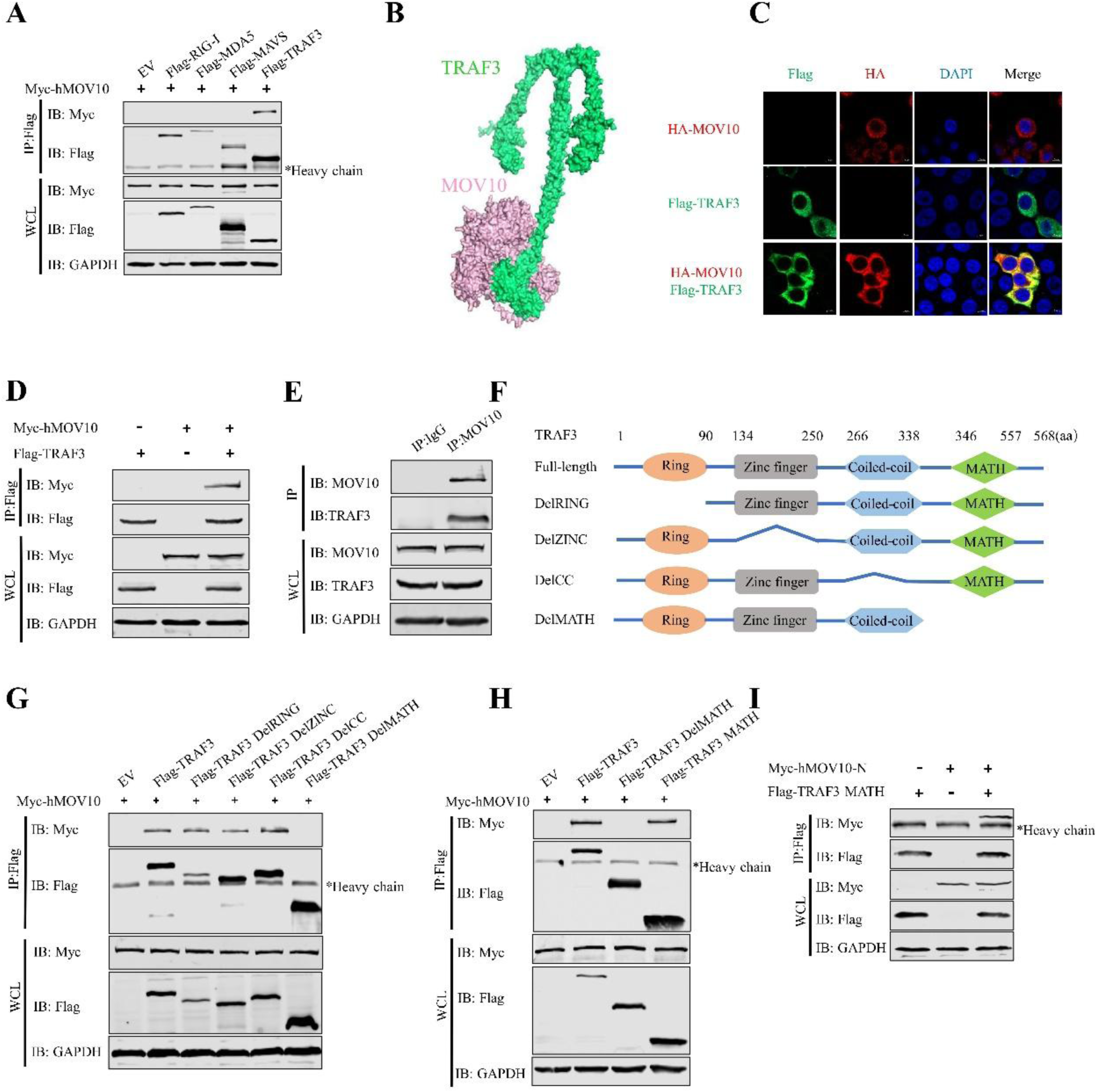
MOV10 interacted with TRAF3 in the RLRs signaling pathway. (A) Plasmids encoding human RIG-I, MDA5, MAVS, and TRAF3 containing a Flag tag were co-transfected into HEK293T cells with Myc-hMOV10, respectively. Cell lysates were harvested and subjected to immunoprecipitation using Flag Ab. The immunoprecipitants were detected by western blotting using Abs against Myc, Flag and GAPDH. (B) Cartoon representation of TRAF3 (green) bound to MOV10 (pink). (C) Co-localization of MOV10 and TRAF3. Vero E6 cells co-transfected with plasmids expressing HA-MOV10 (red) and Flag-TRAF3 (green). Merged images show co-localization of these proteins. Nuclei are highlighted by DAPI staining (blue). (Bar: 5 µm). (D) Exogenous MOV10 interacted with TRAF3. HEK293T cells co-transfected with plasmids for Flag-TRAF3 and Myc-hMOV10, incubated for 24 h, and then collected for Co-IP assay using Flag Ab. (E) Endogenous MOV10 interacted with TRAF3. HEK293T cells were harvested for Co-IP using MOV10 Ab, with IgG Ab as the negative control. Immunoprecipitants were analyzed by western blotting using Abs against MOV10, TRAF3, and GAPDH. (F) Schematic of full-length TRAF3 and its serial truncated mutants. (G and H) Identification of the domain of TRAF3 required for interaction with MOV10. Plasmids encoding TRAF3, DelRING mutant, DelZINC mutant, DelCC mutant, and DelMATH mutant containing Flag tag were co-transfected into HEK293T cells with Myc-hMOV10, respectively. Lysates were harvested and subjected to immunoprecipitation using Flag Ab (G). Plasmids encoding TRAF3, DelMATH mutant, and MATH mutant containing Flag tag were co-transfected into HEK293T cells with Myc-hMOV10, respectively. Co-IP was performed using Flag Ab (H). (I) The N-terminus of MOV10 interacted with the MATH domain of TRAF3. HEK293T cells co-transfected with plasmids for Flag-TRAF3 MATH and Myc-hMOV10-N, incubated for 24 h, and then collected for Co-IP using Flag Ab.

TRAF3 is a member of the tumor necrosis factor receptor associated factors and exhibits RING finger E3 ubiquitin ligase activity[42]. Previous studies have demonstrated that TRAF3 promotes production of IFN-I by interacting with TBK1 and IKKε, playing a critical regulatory role in antiviral immune responses[43, 44]. It is composed of RING finger domain (residues 68-77), zinc finger domain (residues 135-249), coiled coil region domain (residues 267-338), and MATH domain (residues 415-560) [45]. To identify the critical domain of TRAF3 involved in its interaction with MOV10, we constructed several TRAF3 deletion mutants (Fig. 5F). Their interactions were evaluated using Co-IP. We found that deletion of the C-terminal MATH domain of TRAF3 led to loss of interaction with MOV10 (Fig 5G-I). Collectively, these results suggest that MOV10 modulates TRAF3-dependent signaling in the RIG-I-mediated innate immune pathway.

### MOV10 N-terminal domain enhances K63-linked polyubiquitination of TRAF3

TRAF3 is a crucial signaling adaptor protein whose function is finely regulated by ubiquitination[46]. To investigate whether MOV10 modulates ubiquitination of TRAF3, plasmids encoding MOV10, Ub, and TRAF3 were co-transfected into HEK293T cells. We observed that MOV10 enhanced ubiquitination level of TRAF3 (Fig. 6A). Given that K63-linked ubiquitination of TRAF3 is required for induction of IFN-I[42, 46], we next examined the type of ubiquitin linkage in TRAF3 that was promoted by MOV10. After transfection K48- or K63-specific ubiquitin linkage mutants, we found that MOV10 preferentially enhanced K63-linked, rather than K48-linked, polyubiquitination of TRAF3 (Fig. 6B). To further exclude the possibility that MOV10 played a role in regulating the K63-linked polyubiquitination of TRAF3, we examined the stability of TRAF3.

**Figure 6.**
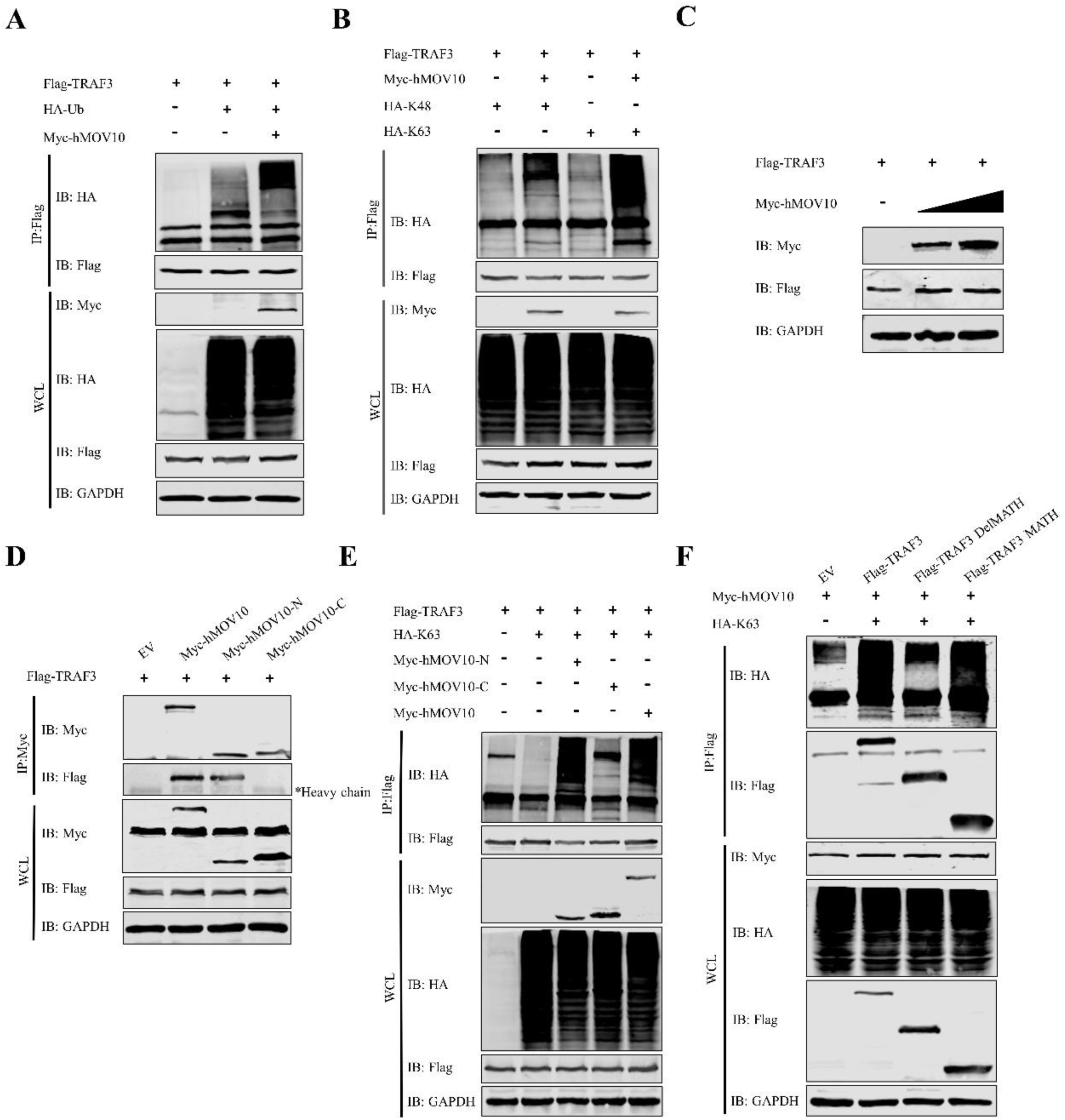
MOV10 enhanced K63-linked polyubiquitination of TRAF3. (A) HEK293T cells were transfected or co-transfected with plasmids encoding Flag-TRAF3, HA-ubiquitin (Ub), and Myc-hMOV10. Cell lysates were collected and subjected to immunoprecipitation using Flag Ab. (B) HEK293T cells were co-transfected with plasmids encoding Flag-TRAF3 and Myc-hMOV10, HA-ubiquitin (K48) or HA-ubiquitin (K63). Lysates were collected and subjected to immunoprecipitation using Flag Ab. (C) HEK293T cells were transfected with plasmids encoding Flag-TRAF3 and Myc-hMOV10 (0, 1, and 3 μg). Lysates were analyzed by western blotting. (D) HEK293T cells were co-transfected with plasmids encoding Flag-TRAF3, full-length Myc-hMOV10 or its mutants (Myc-hMOV10-N and Myc-hMOV10-C). Lysates were collected and subjected to immunoprecipitation using Myc Ab. (E) HEK293T cells were transfected or co-transfected with plasmids encoding Flag-TRAF3, HA-K63, Myc-hMOV10-N, Myc-hMOV10-C, and Myc-hMOV10. Lysates were collected and then subjected to immunoprecipitation using Flag Ab. (F) HEK293T cells were co-transfected with plasmids encoding Myc-hMOV10 and HA-K63, along with plasmids encoding Flag-TRAF3, Flag-TRAF3 DelMATH, or Flag-TRAF3 MATH. Lysates were collected and then subjected to immunoprecipitation using Flag Ab.

Overexpression of increasing amounts of MOV10 in HEK293T cells did not significantly affect the protein level of TARF3 (Fig. 6C). Considering that MOV10 suppressed SADS-CoV replication through the N-terminal domain; therefore, it was interesting to explore whether the N-terminal domain interacted with TRAF3 and enhanced its ubiquitination level. Results shown that the N-terminal domain of MOV10 was crucial for its interaction with TRAF3 (Fig. 6D). Moreover, the deletion mutant lacking the N-terminal domain failed to promote TRAF3 ubiquitination (Fig. 6E). Additionally, the MATH domain was essential for MOV10 to enhance the K63-linked polyubiquitination of TRAF3 (Fig. 6F). In conclusion, these data in combination indicated that MOV10 enhances that K63-linked polyubiquitination of TRAF3 and the antiviral innate immune response, independent of its helicase activity.

### TRIM24 mediates MOV10-enhanced K63-linked polyubiquitination of TRAF3

Since MOV10 not an E3 ligase, we hypothesized that MOV10 likely recruits an E3 ligase for promotion of K63-linked polyubiquitination of TRAF3. A recent study has shown that viral infection induced TRIM24 translocation from the nuclei to the mitochondria and mediated K63-linked ubiquitination of TRAF3[47]. Therefore, we investigated whether the effect of MOV10 on the ubiquitination of TRAF3 was dependent on TRIM24. We first assessed the interaction between MOV10 and TRIM24. Co-IP revealed that MOV10 interacted with TRIM24 (Fig. 7A). Confocal microscopy examined the co-localization of MOV10 and TRIM24 in Vero E6 cells. We observed that MOV10 was distributed throughout the cytoplasm, while TRIM24 was predominantly localized in the nuclei. Notably, co-transfection of MOV10 and TRIM24 led to a significant increase of TRIM24 in the cytoplasm, where it co-localized with MOV10. Furthermore, SeV infection also induced extensive co-localization of TRIM24 with MOV10 (Fig. 7B). Immunoblot analysis also indicated that MOV10 overexpression and SeV infection reduced nuclear TRIM24 levels while increasing its cytoplasmic levels (Fig. 7C). These findings revealed that TRIM24 translocates to the cytoplasm to regulate antiviral immune signaling upon MOV10 transfection or SeV infection.

**Figure 7.**
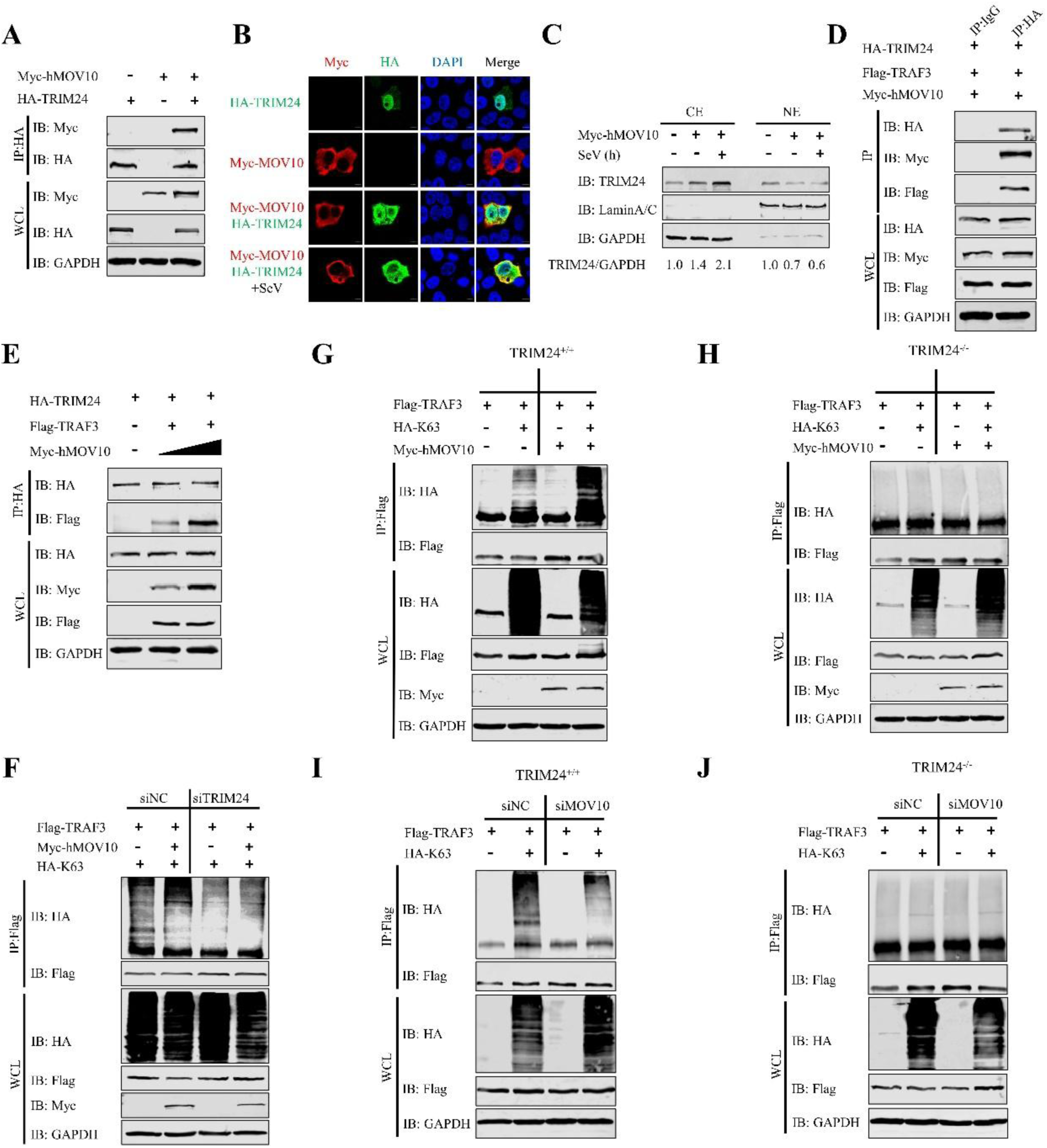
MOV10 promoted TRAF3 K63-linked polyubiquitination dependent on E3 ligase TRIM24. (A) HEK293T cells were co-transfected with Myc-hMOV10 and HA-TRIM24 for 24 h. Cell lysates were harvested and subjected to immunoprecipitation using HA Ab. (B) Co-localization of MOV10 and TRIM24. Plasmids expressing Myc-MOV10 (red) and HA-TRIM24 (green) were co-transfected into Vero E6 cells, which were then infected or not with SeV for 9 h. Merged images show co-localization of these proteins. Nuclei are highlighted by DAPI staining (blue). (Bar: 5 µm). (C) HEK293T cells were transfected with plasmids expressing Myc-hMOV10 or EV and infected or not with SeV. Protein expression of TRIM24, LaminA/C, and GAPDH in cytoplasmic (CE) and nuclear (NE) fractions was assessed by western blotting. The density of TRIM24 bands relative to GAPDH was calculated using grayscale analysis. (D) HEK293T cells were co-transfected with plasmids expressing HA-TRIM24, Flag-TRAF3, and Myc-hMOV10. Cells were harvested for Co-IP using HA Ab, with IgG Ab as the negative control. (E) Plasmids encoding HA-TRIM24, Flag-TRAF3, or Myc-hMOV10 (0, 1 or 3 μg) were co-transfected into HEK293T cells. Cell lysates were collected and then subjected to immunoprecipitation using HA Ab. (F) HEK293T cells were transfected with either siTRIM24 or siNC (50 nM) for 36 h, followed by co-transfection with Flag-TRAF3, HA-K63, or Myc-hMOV10. Lysates were subjected to immunoprecipitation using Flag Ab. (G and H) WT (TRIM24^+/+^) and TRIM24 knockout (TRIM24^−/−^) HEK293T cells were co-transfected with Flag-TRAF3, HA-K63, or Myc-hMOV10. Lysates were subjected to immunoprecipitation using Flag Ab. (I and J) TRIM24^+/+^ and TRIM24^−/−^ HEK293T cells were transfected with either siMOV10 or siNC, followed by co-transfection with Flag-TRAF3 and HA-K63. Lysates were subjected to immunoprecipitation using Flag Ab.

To further confirm the function of MOV10 in promoting TRIM24-dependent TRAF3 K63-linked polyubiquitination, we focused on the interaction between MOV10, TRIM24, and TRAF3. As expected, Co-IP revealed an interaction between MOV10, TRIM24, and TRAF3 (Fig. 7D). Interestingly, we also found that MOV10 enhanced the interaction between TRAF3 and TRIM24 in a dose-dependent manner (Fig. 7E). Subsequently, we investigated the effect of MOV10 on TRAF3 ubiquitination following the knockdown or knockout of TRIM24. We selected siTRIM24-1 with the most significant knockdown effect for subsequent experiments (Extended Data Fig. 2C and D). Compared to the control group, knockdown of TRIM24 in HEK293T cells inhibited enhancement of K63-linked ubiquitination of TRAF3 by MOV10 (Fig. 7F). We then generated TRIM24 knockout (TRIM24^−/−^) HEK293T cells (Extended Data Fig. 3D and E). As shown in Fig. 7G and H, overexpression of MOV10 in WT cells led to enhanced K63-linked polyubiquitination of TRAF3, which was not observed in TRIM24^−/−^ cells. There findings showed that TRIM24 is indispensable in the mediation of TRAF3 K63-linked polyubiquitination by MOV10. To further explore the relationship between MOV10 and TRIM24 in regulating TRAF3 ubiquitination, we knocked down MOV10 in the presence or absence of TRIM24. We observed that knockdown of MOV10 in the presence of TRIM24 reduced K63-linked polyubiquitination of TRAF3. However, knockdown of MOV10 in the absence of TRIM24 did not further decrease TRAF3 ubiquitination (Fig. 7I and J). These results suggested that MOV10 is dependent on TRIM24 in mediating the K63-linked polyubiquitination of TRAF3. In summary, these data suggested that MOV10 serves as a scaffold protein to recruit TRIM24 into mitochondria to promote TRAF3 ubiquitination and aggregation to regulate anti-viral innate immunity.

### SADS-CoV N protein degrades TRAF3 through catalysis of K48-linked polyubiquitination

We found that MOV10 inhibits SADS-CoV replication by modulating TRAF3 to promote IFN production, cell death persists during the later stages of viral infection. Many viruses, particularly RNA viruses, have evolved mechanisms to escape the effects of IFN in the later stages of infection, which prevent effective inhibition of viral replication[21]. To further investigate whether the signaling pathway mediated by MOV10 underwent changes during the later stages of SADS-CoV infection, we measured expression of MOV10, TRAF3, and TRIM24 at different time points in IPI-2I cells infected with SADS-CoV. Notably, we observed that MOV10 expression increased progressively following viral infection, while TRIM24 levels remained unchanged. In contrast, expression of TRAF3 gradually decreased (Fig. 8A). As TRAF3 is a critical regulatory factor in the IFN-I signaling pathway, its reduced expression inhibits IFN production[44]. The mechanism by which SADS-CoV interferes with TRAF3 function warrants further investigation. Li et al. demonstrated that the NS1 protein of influenza A virus impairs the interaction between MOV10 and viral nucleoprotein, which attenuates the antiviral activity of MOV10[48]. Furthermore, NS1 induces MOV10 degradation via the lysosomal pathway[48]. Given that we have identified an interaction between the N protein of SADS-CoV and MOV10, and that expression of TRAF3 was downregulated during viral infection; therefore, we analyzed whether the N protein interacted with TRAF3, which was confirmed by Co-IP (Fig. 8B). We also observed colocalization of these two proteins via confocal microscopy (Fig. 8C). Furthermore, we found that the MATH domain of TRAF3 was indispensable for its interaction with the N2 region of SADS-CoV N protein (Extended Data Fig. 4A-C). These findings indicate that the N protein of SADS-CoV interacts with TRAF3. To further assess the relationship between the viral N protein, MOV10, and TRAF3, we then co-transfected the plasmids encoding these three proteins into HEK293T cells. Notably, Co-IP revealed a clear interaction between MOV10 and TRAF3, which was abolished in the presence of the N protein (Fig. 8D). Similarly, overexpression of the viral N protein disrupted the interaction between endogenous MOV10 and TRAF3 (Fig. 8E). These results proposed that the N protein of SADS-CoV disrupts the interaction between MOV10 and TRAF3.

**Figure 8.**
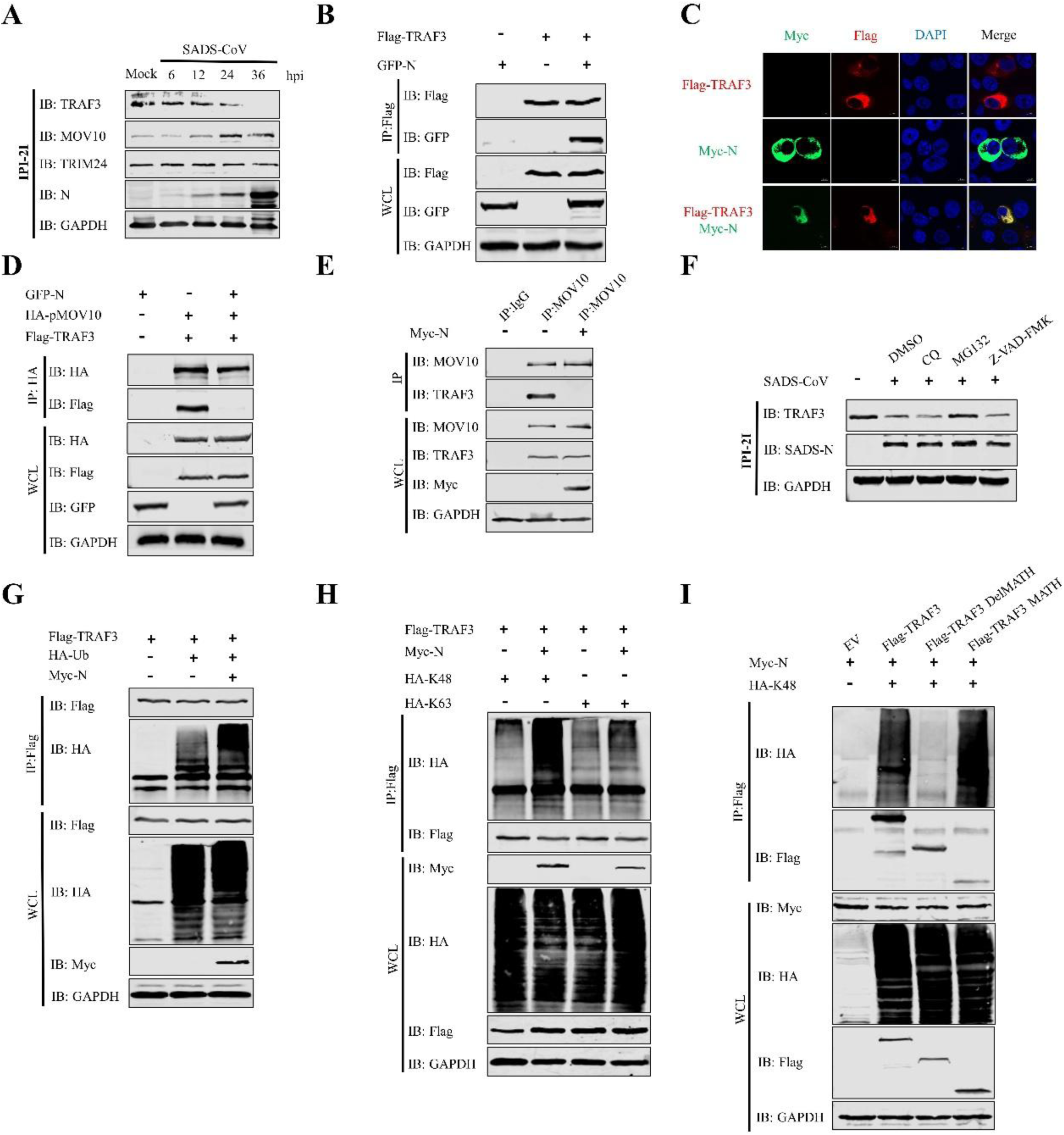
SADS-CoV N protein degraded TRAF3 by catalyzing K48-linked polyubiquitination. (A) IPI-2I cells were infected with SADS-CoV (MOI = 0.1). Cells were collected at 6, 12, 24, and 36 h and detected by western blotting using Abs against TRAF3, MOV10, TRIM24, SADS-CoV N protein and GAPDH. (B) TRAF3 interacted with SADS-CoV N protein. HEK293T cells were co-transfected with Flag-TRAF3 and GFP-N. Cell extracts were subjected to immunoprecipitation using Flag Ab. (C) Co-localization of TRAF3 and SADS-CoV N protein. Plasmids expressing Flag-TRAF3 (red) and Myc-N (green) were transfected or co-transfected into Vero E6 cells. Merged images show co-localization of these proteins. Nuclei are highlighted by DAPI staining (blue). (Bar: 5 µm). (D) Plasmids encoding GFP-N, HA-pMOV10, and Flag-TRAF3 were transfected or co-transfected into HEK293T cells, respectively. Lysates were subjected to immunoprecipitation using HA Ab. (E) HEK293T cells were transfected with plasmids encoding Myc-N or EV. Lysates were subjected to immunoprecipitation using MOV10 Ab, with IgG Ab as the negative control. (F) IPI-2I cells were infected with SADS-CoV (MOI = 0.1) and treated with CQ, MG132, Z-VAD-FMK, or DMSO (all 20 μM) for 5 h before sample collection. Western blotting was used to detect expression of TRAF3, SADS-CoV N protein and GAPDH. (G) HEK293T cells were transfected or co-transfected with plasmids encoding Flag-TRAF3, HA-Ub, and Myc-N. Lysates were collected and then subjected to immunoprecipitation using Flag Ab. (H) HEK293T cells were co-transfected with plasmids encoding Flag-TRAF3, Myc-N, HA-K48 or HA-K63. Lysates were collected and then subjected to immunoprecipitation using Flag Ab. (I) HEK293T cells were co-transfected with Myc-N, HA-K48, Flag-TRAF3 or Flag-TRAF3 MATH. Lysates were subjected to immunoprecipitation using Flag Ab.

Cellular protein degradation is primarily mediated by the ubiquitin-proteasome system and autophagy[49, 50]. Additionally, proteolytic cleavage by caspases is a key feature of apoptosis, a form of programmed cell death[51]. To investigate the pathways by which SADS-CoV degraded TRAF3, cells were treated with CQ (an autophagy inhibitor), MG132 (a ubiquitin-proteasome inhibitor), and Z-VAD-FMK (an apoptosis inhibitor) to measure TRAF3 expression. Cell viability assay showed that none of the three compounds exhibited cytotoxicity in IPI-2I cells at 20 μM (Extended Data Figure 4D). Compared to the DMSO group, TRAF3 expression showed no significant difference after CQ and Z-VAD-FMK treatment, while it was obviously upregulated after MG132 treatment (Fig 8F). This suggested that the SADS-CoV N protein degraded TRAF3 via the ubiquitin proteasome pathway. We next investigated whether the N protein modulated the ubiquitination level of TRAF3. Plasmids encoding SADS-CoV N protein, ubiquitin, and TRAF3 were co-transfected into HEK293T cells. As expected, a significant upregulation of TRAF3 ubiquitination levels was observed in the presence of SADS-CoV N protein (Fig. 8G). Given that different ubiquitin linkages are associated with distinctive functions, we next investigated the types of ubiquitin linkages in TRAF3 induced by the SADS-CoV N protein. Plasmids encoding K48- or K63-specific ubiquitin mutants were co-transfected with TRAF3 and the SADS-CoV N protein into HEK293T cells. We found that the N protein markedly enhanced K48-linked polyubiquitination of TRAF3 (Fig. 8H). Furthermore, the MATH domain of TRAF3 was essential for the SADS-CoV N protein-mediated K48-linked polyubiquitination of TRAF3 (Fig. 8I). Taken together, these results suggest that the SADS-CoV N protein antagonizes the host immune response through two mechanisms: firstly, by disrupting the interaction between MOV10 and TRAF3, and secondly, by catalyzing K48-linked polyubiquitination to promote degradation of TRAF3.

## Disscusion

In this study, we identified that MOV10, a superfamily 1 (SF1) RNA helicase, suppressed SADS-CoV replication by enhancing TRIM24-mediated K63-linked ubiquitination of TRAF3, which promotes IFN production. However, the antiviral activity of MOV10 was antagonized by viral N protein, which disrupted the interaction of MOV10 with TRAF3. Furthermore, the viral N protein directly degraded TRAF3 through K48-linked ubiquitination, inhibiting IFN production during the later stages of SADS-CoV replication. These results shed light on the mechanism by which MOV10 mediates antiviral immunity and demonstrate that SADS-CoV evades the host’s immune defenses (Fig. 9).

**Figure 9.**
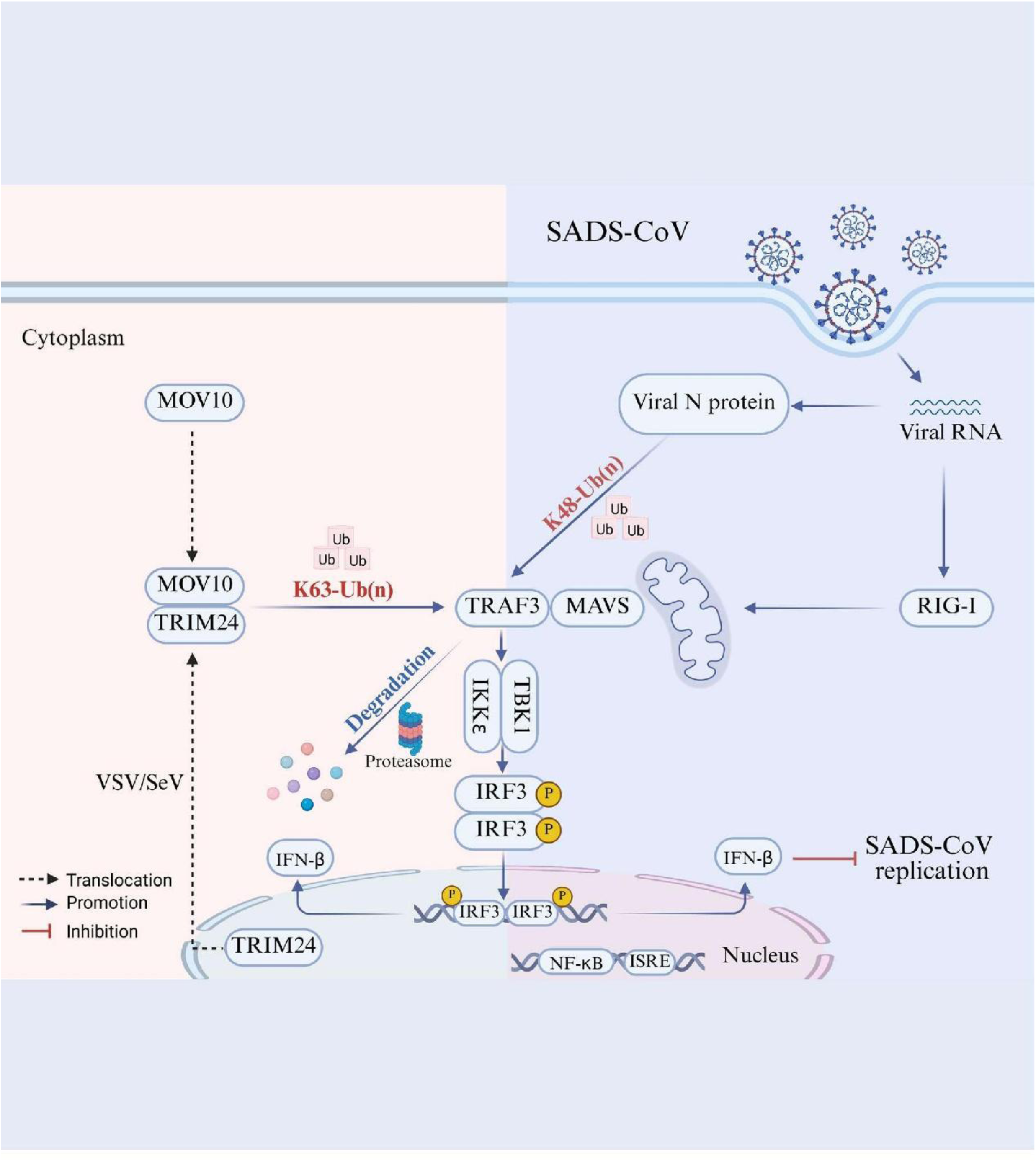
Models of MOV10 mechanisms in counteracting SADS-CoV and viral N protein antagonizing host innate immunity. To inhibit SADS-CoV replication, the host factor MOV10 enhanced the catalysis of K63-linked polyubiquitination of TRAF3 by E3 ubiquitin ligase TRIM24, thereby activating the RIG-I-mediated antiviral signaling pathway. On the other hand, the SADS-CoV N protein suppressed RIG-I-mediated antiviral signaling by promoting K48-linked polyubiquitination, leading to degradation of TRAF3.

MOV10 has been reported to inhibit the replication of various viruses through distinct mechanisms. For instance, it suppresses influenza A virus replication by blocking the nuclear import of the viral nucleoprotein[48, 52]. In bunyaviruses, MOV10 binds to the N-arm domain of the nucleoprotein, thereby preventing N polymerization and ribonucleoprotein (RNP) assembly[53]. In HIV-1, MOV10 exerts antiviral effects by enhancing nuclear export of viral mRNA, interacting with the Gag protein, and being incorporated into viral particles, with its C-terminal region and helicase motifs being essential for this activity. Its interaction with Rev further facilitates nuclear export of viral RNA[54–56]. Additionally, MOV10 restricts reverse transcription in viruses such as hepatitis B[57] virus and HIV[54]. In the context of coronaviruses, MOV10 inhibits MERS-CoV and SARS-CoV-2 by interacting with the viral nucleocapsid protein and leveraging its helicase activity to suppress viral RNA replication[35, 58]. MOV10 also exerts antiviral effects by promoting IFN production, as observed in vesicular stomatitis virus (VSV)[41] and herpes simplex virus type 1[59]. In the present study, we demonstrated that MOV10 inhibited SADS-CoV replication by upregulating IFN-β expression, which was mediated by the N-terminal domain of MOV10. However, the molecular mechanism for the antiviral activity of MOV10 varies among coronaviruses. In contrast to SADS-CoV, MERS-CoV exhibits a divergent requirement for MOV10, where the helicase activity is indispensable while the N-terminal domain is dispensable[35]. Furthermore, it remains unclear whether the helicase activity is necessary for its action against SARS-CoV-2[58]. These differences suggest that MOV10 may exerts broad antiviral effects within the coronavirus family through multiple mechanisms.

The production of IFN-I, initiated by recognition of pattern recognition receptors (PRRs), plays a crucial role in the innate immune defense against viral infections. Our recent study showed that SADS-CoV infection activates the RIG-I and MAVS signaling pathways, while host protein DDX11 promotes activation of this signaling pathway through its interaction with RIG-I and MAVS. This activation cascade induces phosphorylation of TBK1 and IRF3, which subsequently induces IFN production[29]. However, regarding the role of MOV10 in innate immunity, Cuevas et al. demonstrated that it restricts VSV proliferation by promoting IFN production via enhancement of IKKε, independently of TBK1 and the RIG-I-MAVS pathway[41]. Furthermore, this conclusion is strengthened by the finding that MOV10 also inhibits HSV infection by promoting IFN production via an IKKε-mediated RNA-sensing pathway, independently of RIG-I, MDA-5, and MAVS[59]. This suggests that MOV10 inhibits viral infection by inducing production of IFN through a unique mechanism. Here, we identified a novel role for MOV10 in TRAF3 ubiquitination, a mechanism not previously reported. Specifically, the N-terminal region of MOV10 enhanced K63-linked ubiquitination of TRAF3, with the MATH domain of TRAF3 playing a crucial role in this process. The function of TRAF3 in signaling is intricately linked to its ubiquitination status; a crucial post-translational modification recognized as a key regulator of host IFN-I signaling pathways[60, 61]. TRAF3 undergoes two distinct types of ubiquitination: K48-linked polyubiquitination, which targets it for proteasomal degradation; and K63-linked polyubiquitination, which does not lead to degradation but likely modulates its function in signaling pathways[42, 46]. Critically, K63-linked polyubiquitination of TRAF3 is essential for the recruitment and activation of TBK1/IKKε. This activation triggers the phosphorylation of the transcription factors IRF3/7, ultimately driving induction of IFN-I[47, 62]. Therefore, by promoting K63-linked polyubiquitination of TRAF3, MOV10 facilitates IFN production, which in turn inhibits SADS-CoV replication.

MOV10 has not been reported to have E3 ubiquitin ligase activity. Therefore, it is likely that other E3 ligases are involved in ubiquitination of TRAF3, with MOV10 facilitating the process indirectly, possibly by interacting with these ligases or modifying their activity. Zhu et al. recently discovered that TRIM24 is a direct E3 ligase, which specifically induces K63-linked ubiquitination of TRAF3 during SeV/VSV infection[47]. They found that RNA virus infection triggers CRM1-dependent TRIM24 translocation from the nuclei to the mitochondria, where it ubiquitinates TRAF3, activating the antiviral transcription factor IRF3, which then initiates IFN-I gene transcription. Consistent with these findings, we demonstrated that upon co-transfection of MOV10 and TRIM24, TRIM24 was also translocated from the nuclei to the cytoplasm, where it co-localized with MOV10. Furthermore, an interaction between MOV10, TRIM24, and TRAF3 was identified. Notably, our data indicated that TRIM24 promotes K63-linked polyubiquitination of TRAF3, with MOV10 enhancing the efficiency of this process. Our identification of the role of TRIM24 reinforces the critical importance of E3-ligase-mediated ubiquitination in controlling TRAF3 activity; a mechanism shared by several other enzymes including TRIM35, HECTD3, Nedd4l, RNF166, and RNF126, which have been identified as key regulators of immune signaling pathways[62–66]. These enzymes regulate activation of IFN-I by mediating ubiquitination of TRAF3, thereby enhancing the host immune response against pathogens.

Viruses have evolved elaborate mechanisms to evade or inactivate the innate immune signaling pathway for their replication[67]. Ubiquitin modifications and interplay between proteins in these signaling pathways enhance IFN-I expression. Conversely, viruses exploit the ubiquitin modification system to inhibit IFN-I production[68, 69]. In addition to illustrating the anti-SADS-CoV activity and mechanism of MOV10, we also uncovered a viral counterstrategy mediated by the N protein of SADS-CoV. Mechanically, we found that the N protein inhibited the antiviral effect of MOV10 by disrupting the interaction between MOV10 and TRAF3. Moreover, the N protein promoted K48-linked polyubiquitination of TRAF3, leading to TRAF3 degradation via the ubiquitin-proteasome pathway. Through these two mechanisms, SADS-CoV effectively suppresses IFN production. Actually, the N protein of coronaviruses plays a key role in suppressing the secretion of host IFN-I, which facilitates viral evasion of the host immune surveillance system. Several viruses, including SARS-CoV-2[10], SARS-CoV[11], PEDV[14], PDCoV[13], and MHV[12] inhibit IFN-β production through their N proteins using different mechanisms. Notably, Zhou et al. revealed that SADS-CoV antagonizes IFN-β production via blocking the interaction between TRAF3 and TBK1[15]. Building on this, our study further elucidated the molecular mechanism by which the SADS-CoV N protein promotes degradation of TRAF3 through the ubiquitin-proteasome pathway. This degradation likely impairs the ability of TRAF3 to interact with downstream signaling molecules, ultimately inhibiting production of IFN. Therefore, while Zhou et al. highlighted the disruption of TRAF3-TBK1 interaction as a key mechanism, our findings provide additional insights into how SADS-CoV N protein facilitates immune evasion by directly targeting TRAF3 for degradation.

In summary, we have revealed a novel mechanism by which MOV10 antagonizes SADS-CoV. MOV10 suppresses SADS-CoV replication by promoting TRIM24-mediated K63-linked ubiquitination of TRAF3, which in turn enhances IFN production. However, we also found that the N protein of SADS-CoV counters the antiviral effect of MOV10. The N protein not only inhibits the antiviral effect of MOV10 by disrupting the interaction between MOV10 and TRAF3, but also promotes K48-linked polyubiquitination of TRAF3, leading to TRAF3 degradation via the ubiquitin-proteasome pathway. These findings highlight the complex interplay between MOV10 and SADS-CoV, suggesting that MOV10 may functions as a broad-spectrum antiviral agent. Further research is needed to fully understand its role across different viral infections. Collectively, these insights could inform the development of novel antiviral strategies, particularly those targeting SADS-CoV infection.

## Materials and Methods

### Cells and viruses

Porcine intestinal epithelial cells (IPI-2I), swine testis cells (ST), African green monkey kidney cells (Vero E6), and human embryonic kidney cells (HEK293T) were cultured in Dulbecco’s Modified Eagle’s Medium (DMEM; 11965092; Life Technologies, Carlsbad, CA, USA)) supplemented with 100 mg/mL streptomycin, 100 units/mL penicillin, and 10% fetal bovine serum (10099141C; Gibco, Life Technologies). The cells were maintained at 37°C in a 5% CO_2_ incubator. SADS-CoV (GenBank accession number: MF094681) was described previously[25], as was Sendai virus (SeV)[26].

### Antibodies and reagents

SADS-CoV N protein specific monoclonal antibody (mAb) 3E9 was prepared and maintained in our laboratory[27]. Abs against Flag (ab205606), hemagglutinin (HA, ab9110), phosphorylated (p)-IRF3 (ab76493), IRF3 (ab68481), and GFP (ab290) were purchased from Abcam (Cambridge, MA, USA). Abs against Myc (M4439), HA (H9658), Flag (F1804), and glyceraldehyde 3-phosphate dehydrogenase (GAPDH, G9545) were purchased from Sigma-Aldrich (St. Louis, MO, USA). Abs against MOV10 (10370-1-AP), TRAF3 (18099-1-AP), TRIM24 (14208-1-AP) and HA (51064-2-AP) were purchased from Proteintech (Wuhan, China). Ab against Lamin A/C was purchased from Beyotime (Shanghai, China). IRDye 800CW goat anti-mouse IgG secondary Ab (926-32210) and IRDye 680RD goat anti-rabbit IgG secondary Ab (925-68071) were purchased from LiCor BioSciences (Lincoln, NE, USA). RNase A, Alexa Fluor 594 goat anti-rabbit IgG (H+L), Alexa Fluor 488 goat anti-mouse IgG (H+L), and beads were purchased from Thermo Fisher Scientific (Waltham, MA, USA). Poly(I:C) was purchased from InvivoGen (Hong Kong, China) and was used at a final concentration of 10 µg/mL. Recombinant human IFN-β and swine IFN-β were purchased from R&D Systems (Minneapolis, MN, USA). MG132, Z-VAD-FMK and CQ were purchased from MedChemExpress (Edison, NJ, USA). The Dual-Luciferase Reporter Assay System was purchased from Promega (Madison, WI, USA).

### Plasmids and transfection

Porcine full-length MOV10 (GenBank accession number: XM_021090739.1) and human full-length MOV10 (GenBank accession number: NM_001130079.3) were amplified from the cDNA of IPI-2I and HEK293T cells, respectively. The DNA fragment of MOV10 was inserted into the pCAGGS-HA (P0166; Miaoling Biology) or pCMV-Myc (635689; Clontech) vector via homologous recombination using the ClonExpress ultra one-step cloning kit (C116; Vazyme, Nanjing, China). The truncated mutants containing of porcine or human MOV10 N-terminal region (MOV10-N_93-305 aa) and C-terminal region (MOV10-C_524-912 aa) were cloned into the pCAGGS-HA or pCMV-Myc vector. SADS-CoV N protein (GenBank accession number: ON911569.1) was cloned into the pAcGFP-C1 (632470; Clontech) and pCMV-Myc vector. The truncated mutants of SADS-CoV N (N1_1-146 aa, N2_147-249 aa, N3_250-376 aa) were constructed on the basis of the full-length SADS-CoV N plasmid and inserted into the pAcGFP1-C1 vector, respectively. The IFN-sensitive response element (ISRE) and IFN-β promoter luciferase reporter plasmid, along with the internal reference plasmid (pRL-CMV) were stored in our laboratory as previously described[28]. Human full-length TRAF3 (GenBank accession number: NM_003300.4) was cloned into pCMV-Flag vector (635688; Clontech). The TRAF3 truncated mutants (DelRING_90-568 aa, DelZINC_1-134 aa and 250-568 aa, DelCC_1-266 aa and 338-568 aa, DelMATH_1-346 aa, MATH_346-568 aa) were amplified from the full-length TRAF3 and inserted into the pCMV-Flag vector to generate the recombinant plasmids Flag-TRAF3-DelRING, Flag-TRAF3-DelZINC, Flag-TRAF3-DelCC, Flag-TRAF3-DelMATH, and Flag-TRAF3-MATH, respectively. The plasmids for expression of Flag-RIG-I, Flag-MDA5, and Flag-MAVS were preserved at our laboratory as previously described[29]. The plasmids HA-ubiquitin (HA-Ub), HA-ubiquitin (K48) and HA-ubiquitin (K63) were stored in our laboratory. HEK293T and IPI-2I cells were transfected with the above plasmids using Lipofectamine^TM^ 3000 transfection reagent (L3000001; Thermo Fisher Scientific).

### Screening of host factors associated with SADS-CoV N protein

Vero E6 cells were infected with SADS-CoV (MOI=1) and incubated for 36 h. Cells were then collected and lysed. The resulting lysates were subjected to immunoprecipitation using Ab 3E9 targeting the N protein of SADS-CoV (1:100 dilution)[27], with IgG at the same concentration as a negative control. Enriched protein complexes were subsequently analyzed using LC-MS/MS to identify host factors potentially interacting with the viral N protein.

### Western blotting

Western blotting was performed as described previously[30]. Cells were lysed on ice for 30 min using radio-immunoprecipitation assay buffer (R0278; Sigma-Aldrich, USA) supplemented with 1 mM phenylmethylsulfonyl fluoride (PMSF) solution (ST506-2; Beyotime). The lysate was then centrifuged at 12000×g for 5 minutes at 4°C to remove cellular debris. Sodium dodecyl sulfate polyacrylamide gel electrophoresis (SDS-PAGE) loading buffer (P0015L; Beyotime) was added to the supernatant and boiled for 10 min. Equal amounts of total proteins were analyzed with 12.5% SDS-PAGE and transferred to nitrocellulose membranes (66485; Pall Corporation, Port Washington, NY, USA). After blocking with 5% skim milk for 1 h at room temperature (RT), the nitrocellulose membranes were incubated with primary Abs for 6-8 h at 4°C, followed by incubation with IRDye 800CW goat anti-mouse lgG (H+L) secondary Abs (1:10000) (926-32210; LiCor BioSciences, USA) or IRDye 680RD goat anti-rabbit lgG (H+L) secondary Abs (1:10000) (926-68071; LiCor BioSciences, USA) for 45 min in the dark. An Odyssey infrared imaging system (LiCor BioSciences, USA) was used to visualize the blots.

### Co-immunoprecipitation (Co-IP) assay

Cells were lysed on ice for 30 min in IP lysis buffer (87788, Thermo Fisher Scientific) containing 1 mg/mL protease inhibitor cocktail (04693132001; Roche, Switzerland). The lysates were then incubated overnight at 4°C with primary Abs, followed by 6-8 h incubation with protein A/G magnetic beads at 4°C. After washing the bead-Ab-antigen complexes five times with lysis buffer, the immunoprecipitated proteins were collected for western blotting.

### RNA extraction, reverse transcription, and quantitative real-time polymerase chain reaction (qRT-PCR)

Total RNA was extracted from treated cells using the Simply P Total RNA Extraction Kit (BSC52S1; BioFlux, Brussels, Belgium) and reverse transcribed into cDNA using the PrimeScript^™^ IV 1st Strand cDNA Synthesis Mix (6215A; Takara, Japan). qRT-PCR was conducted using the QuantStudio^®^ 5 Real-Time PCR System (Applied Biosystems, Foster City, CA, USA) with TB Green^®^ Premix Ex Taq^™^ II (RR820A; Takara, Japan). Relative quantitative mRNA levels were calculated using the 2^-ΔΔCt^ method. All experiments were performed at least three times. The sequences of the primers synthesized for qRT-PCR are listed in Table 1.

**Table 1.**
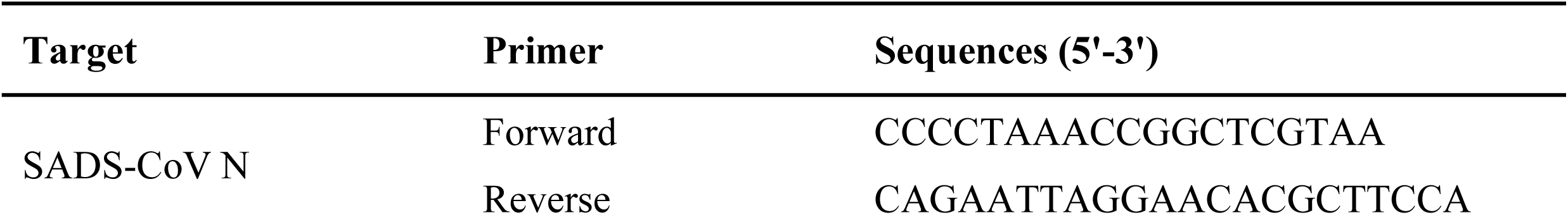
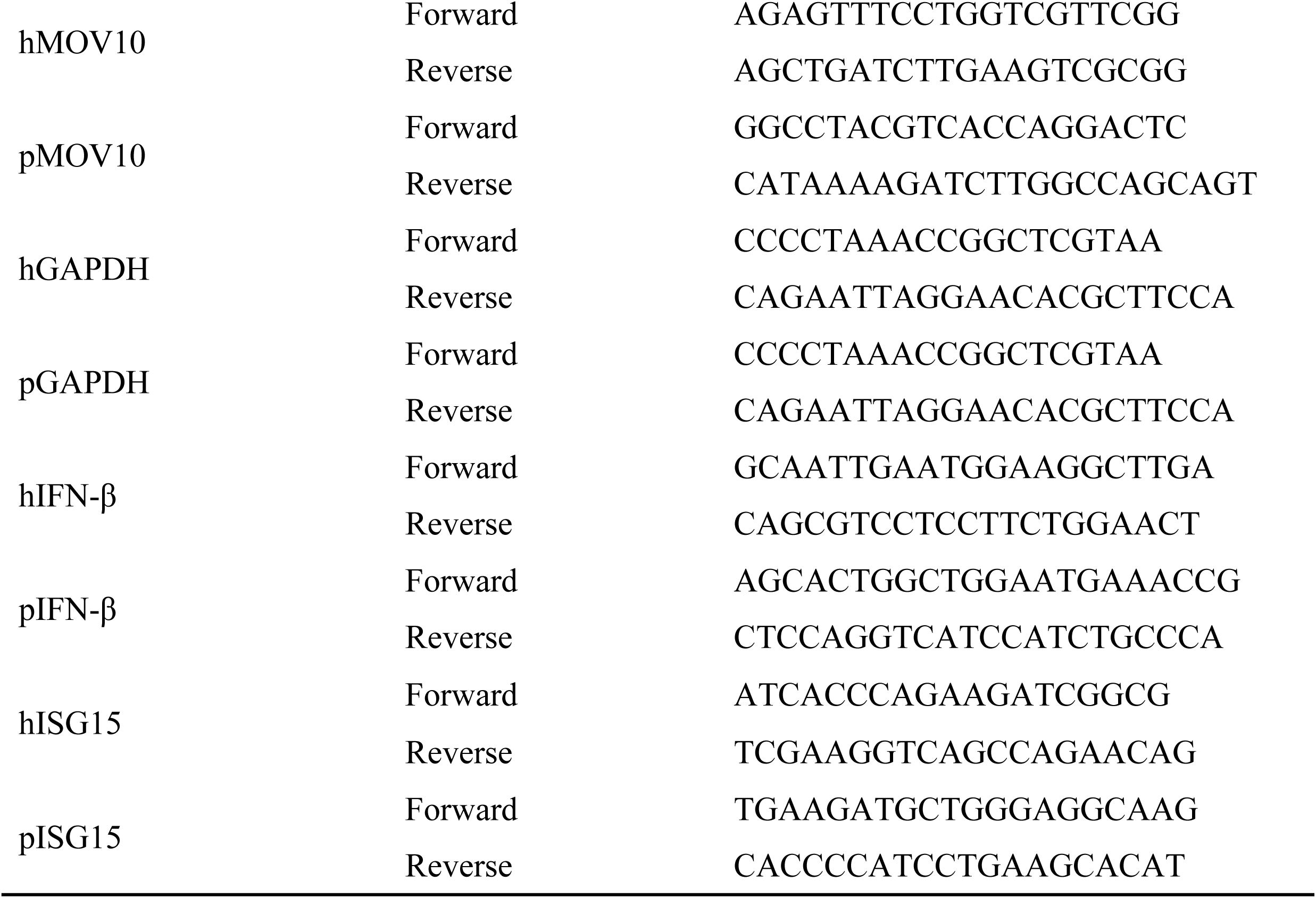
Sense sequences are used for qRT-PCR in this study.

### Animal experiment and immunohistochemistry (IHC) assay

Six 3-day-old specific pathogen-free (SPF) piglets were randomly assigned to either a challenge or control group. The challenge group was orally infected with 5×10^4^ TCID_50_ of SADS-CoV, whereas the control group was orally infected with the same volume DMEM. We recorded the clinical symptoms (vomiting and diarrhea), and euthanized all piglets at 36 hours post-infection (hpi). Intestinal tissues samples were collected for qRT-PCR. Representative sections of the small intestine tissue samples (jejunum, and ileum) were fixed in 4% paraformaldehyde and stored in 70% ethanol at 4°C. IHC was performed as previously described[30]. Slides were incubated overnight at 4°C with mAb 3E9 (1:50) against the SADS-CoV N protein, followed by incubation with horseradish peroxidase-labeled goat anti-mouse IgG_1_ for 1 h. Immunocomplexes were detected using the 3,3ʹ-diaminobenzidine liquid substrate system.

### Construction of knockout cell lines

To generate MOV10^−/−^ IPI-2I cells and TRIM24^−/−^ HEK293T cells, the CRISPR/Cas9 gene-editing system was used. The single-guide RNA (sgRNA) was designed using the website (http://crispor.tefor.net/crispor.py). The specific sequences can be found in Table 2. The sgRNA-1 was cloned into the pX458-EGFP vector. The sgRNA-2 was cloned into the pX458-mCherry vector. Cells were transfected or co-transfected with pX458-sgRNA-1 and pX458-sgRNA-2. After cell culture, positive cells (exhibiting red and green dual fluorescence) were individually sorted into single clones on a 96-well plate using SH800S flow cytometer (Sony, Tokyo, Japan). After expanding and culturing the positive clones, cell samples were collected for sequencing and western blotting.

**Table 2.**
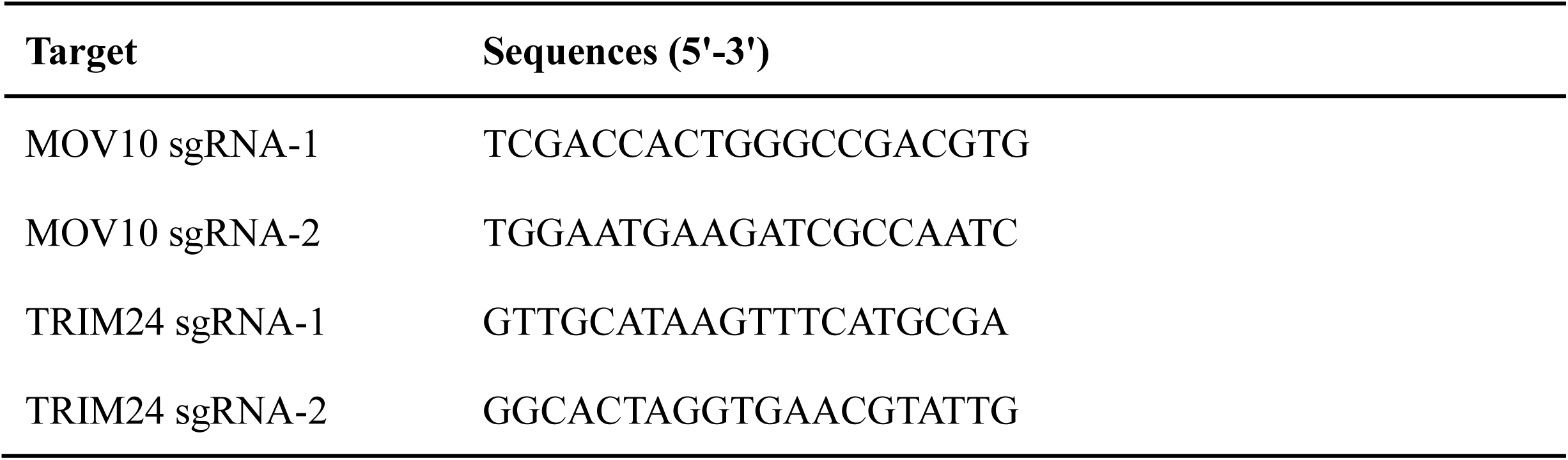
Primer sequences of sgRNAs in this study.

### RNA interference

Small interfering RNAs (siRNAs) targeting porcine MOV10 and human TRIM24 were designed and synthesized by Seven Biotechnology Co., Ltd (Beijing, China). The siRNA target sequences used in the experiments are listed in Table 3. IPI-2I and HEK293T cells were seeded in 12-well plates, grown to 30∼40% confluency, and then transfected for 48 h with 50 nM siRNAs using Lipofectamine RNAiMAX Reagent (13778075; ThermoFisher Scientific). Following transfection, the cells were infected with SADS-CoV (MOI=0.1), and samples were collected 24 hpi.

**Table 3.**
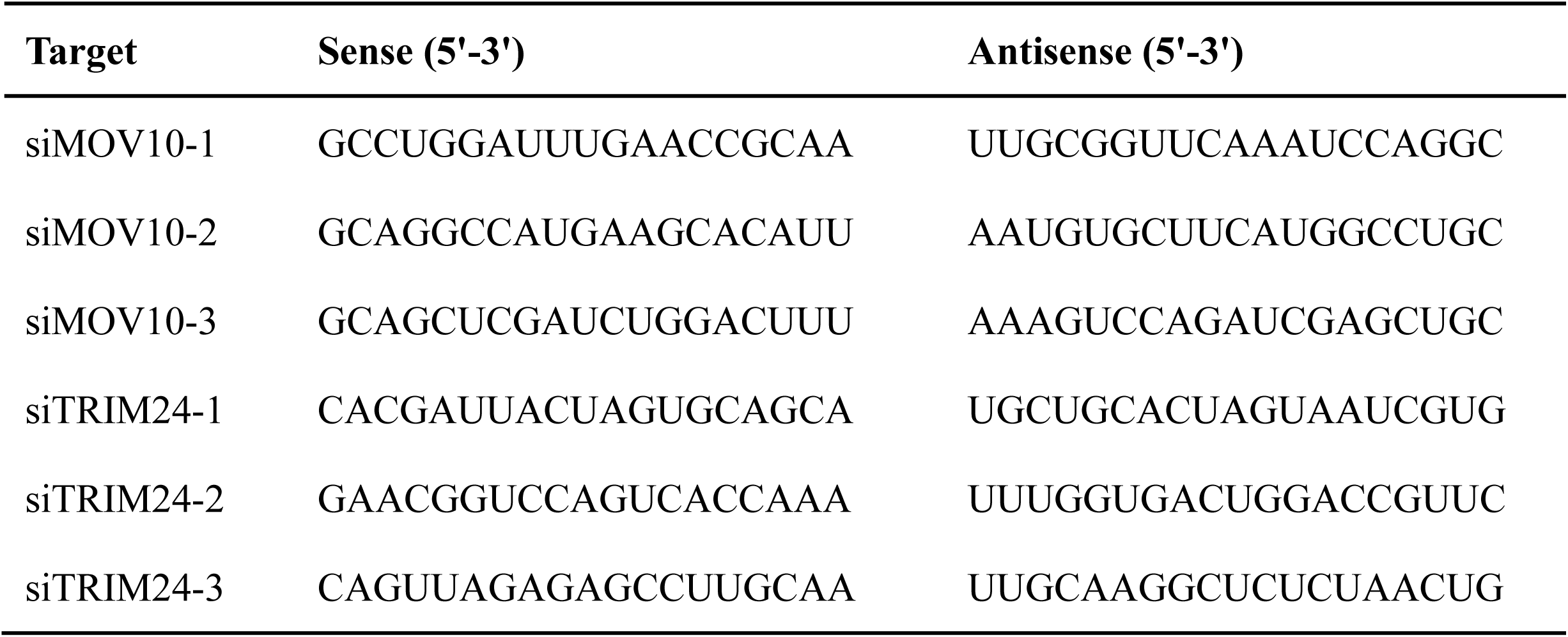
Primer sequences of siRNAs in this study.

### 50% tissue culture infective doses (TCID_50_)

Vero E6 cells were infected with 10-fold serial dilutions of each supernatant. At 4-6 days post-infection, cytopathic effects in cells were observed through microscopy. Viral titers were calculated using the Reed-Muench method.

### Dual-luciferase reporter assays

Dual-luciferase reporter assays were performed as described previously[29]. HEK293T cells were seeded in 24-well plates. The cells were then transfected with either pCMV-Myc-MOV10 or an empty vector (EV), in addition to ISRE-Luc or IFN-β-Luc reporter constructs, along with an internal reference plasmid, pRL-CMV. After 24 hours post transfection (hpt), the cells were infected with SeV for 9 h. Following infection, the cells were lysed, and luciferase activity was assessed using a dual luciferase reporter assay kit (E1901; Promega, Madison, WI, USA).

### Immunofluorescence assay and confocal microscopy

Vero E6 cells cultured in 35-mm diameter dishes were transfected with specific plasmids. Cells were either infected or not infected with SeV after 24 hpt. The cells were fixed with precooled 4% paraformaldehyde (16005; Sigma-Aldrich) at RT for 30 min and permeabilized with 0.1% Triton X-100 (T8787; Sigma-Aldrich) at RT for 15 min. After blocking with 5% skim milk (P0216; Beyotime) at 4°C overnight, the cells were incubated with primary Abs for 1-4 h at RT. After three washes with PBST, the cells were incubated with corresponding fluorescein-conjugated secondary Abs (Alexa Fluor^™^ 594/488) at RT for 1 h in the dark. After PBST washing, cell nuclear were stained with 4’,6-diamidino-2-phenylindole (DAPI) for 15 min and then washed with PBST three times. Finally, the cells were directly observed using an LSM880-ZEISS confocal laser scanning microscope with Fast Airyscan (Zeiss, Germany).

### Cell viability assay

IPI-2I cells were grown in 96-well plates and then treated with drugs for 24 h. Cell viability was determined using Cell Counting Kit-8 (CCK-8; CK04, Dojindo Laboratories Co., Ltd., Kumamoto, Japan) according to the manufacturer’s instructions.

### Ubiquitination assay

To analyze the effect of MOV10 on ubiquitination of TRAF3 in HEK293T cells, HEK293T cells were co-transfected with the indicated plasmids and then whole cell lysates were immunoprecipitated with the corresponding Abs and analyzed by western blotting. To analyze the effect of SADS-CoV N protein on ubiquitination of TRAF3 in transfected cells, HEK293T cells were co-transfected with the indicated plasmids and then whole cell lysates were immunoprecipitated with the corresponding Abs and analyzed by western blotting.

### Statistical analysis

All results shown in the figures were presented, where appropriate, as mean and standard deviation (SD). The results of three independent experiments were analyzed with GraphPad Prism software (version 9.0; San Diego, CA, USA). Error bars represent SD. *P*-values are \**P* < 0.05; \*\**P* < 0.01; \*\*\**P* < 0.001; \*\*\*\**P* < 0.0001; ns, not significant.

## Supporting information

**S1 Data.**
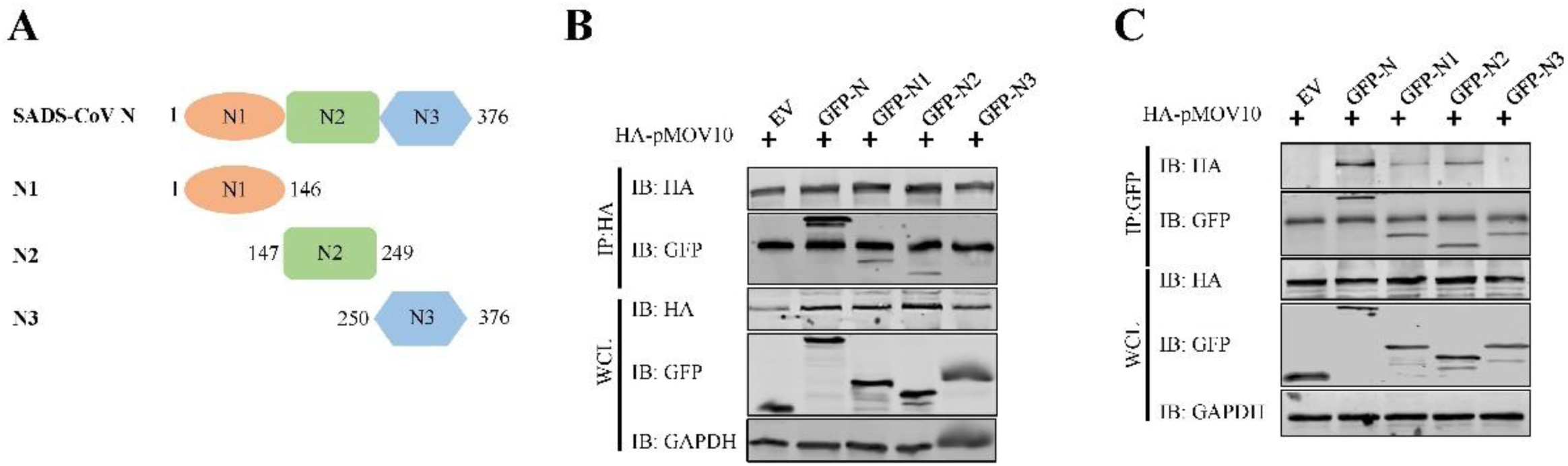
MOV10 interacted with the N1 and N2 regions of SADS-CoV N protein. (A) Schematic of full-length SADS-CoV N and its serial truncated mutants. (B-C) MOV10 interacted with N1 and N2 domains of SADS-CoV N protein. HEK293T cells were transfected or co-transfected with plasmids encoding HA-pMOV10, GFP-N, GFP-N1, GFP-N2, or GFP-N3. Cellular lysates were subjected to immunoprecipitation using HA (B) or GFP (C) Abs.

**S2 Data.**
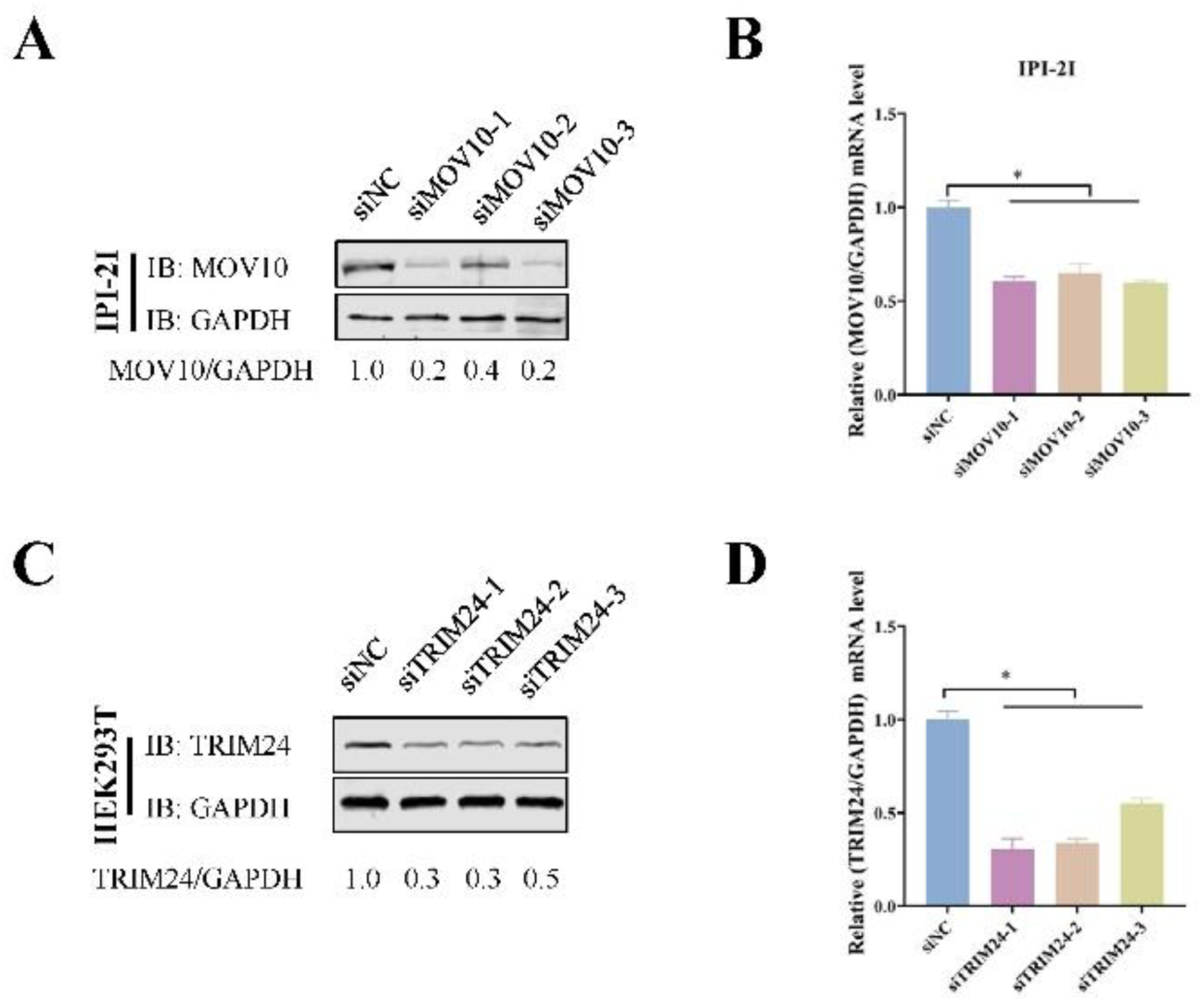
Evaluation of the knockdown efficiency of siRNA targeting MOV10 and TRIM24. (A and B) siRNA-mediated MOV10 knockdown experiment. IPI-2I cells were transfected with siRNAs (siMOV10-1, siMOV10-2 and siMOV10-3) or siNC at 50 nM for 48 h. Western blotting (A) and qRT-PCR (B) assessed the MOV10 knockdown efficiency. Band density for MOV10/GAPDH was calculated, with values from the siNC group standardized to 1. (C and D) siRNA-mediated TRIM24 knockdown experiment. HEK293T cells were transfected with three siRNAs against porcine TRIM24 and siNC at 50 nM for 48 h. Western blotting (C) and qRT-PCR (D) assessed expression of TRIM24. Band density for MOV10/GAPDH was calculated, with values from the siNC group standardized to 1. The mean and SD of the results from three independent experiments are shown (\**P* < 0.05).

**S3 Data.**
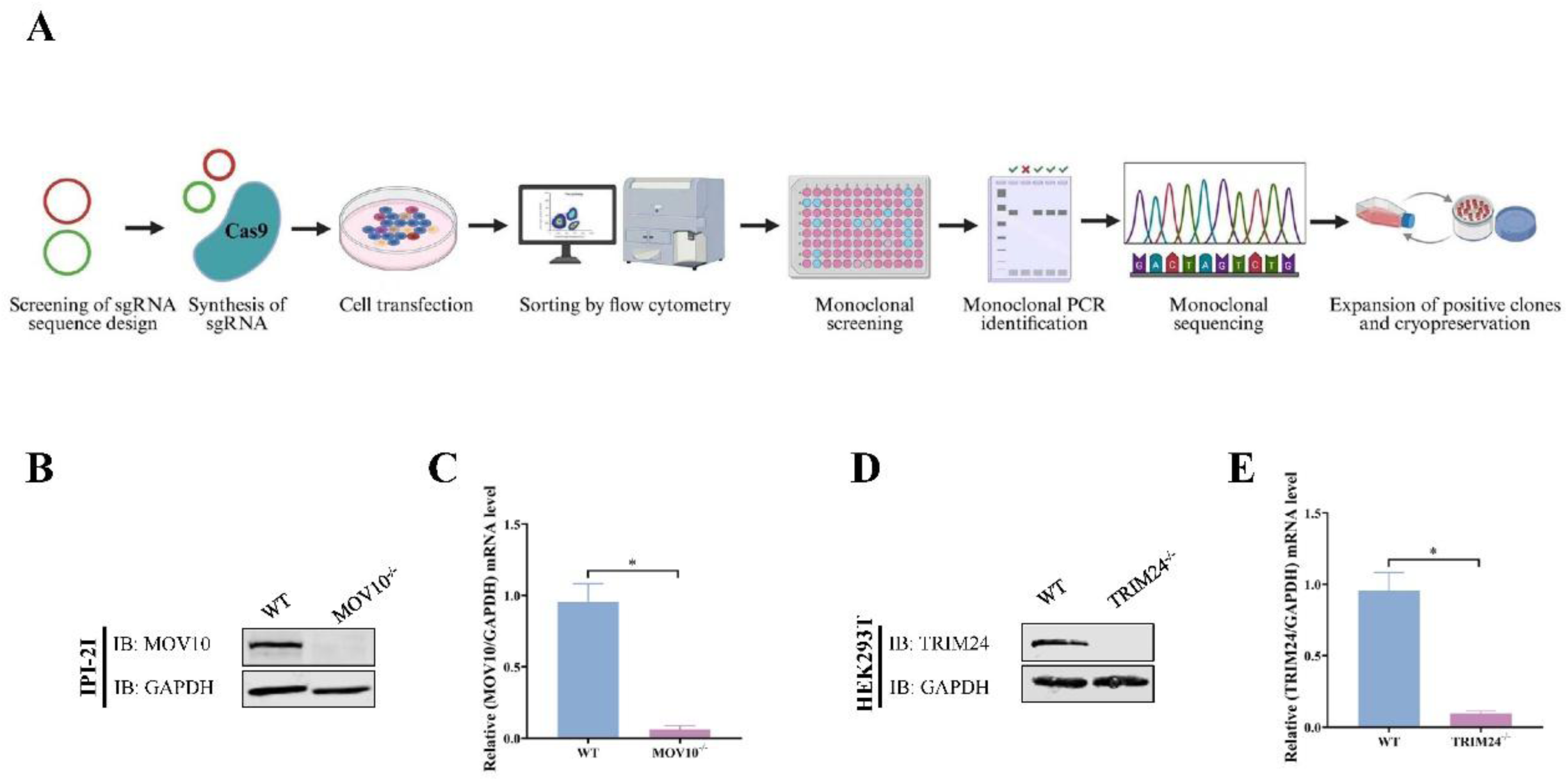
Generation MOV10^−/−^ IPI-2I cells and TRIM24^−/−^ HEK293T cells. (A) Schematic of CRISPR/Cas9-based knockout cell line construction. (B and C) Expression of MOV10 in WT and MOV10^−/−^ IPI-2I cells were assessed by western blotting and qRT-PCR. (D and E) Expression of TRIM24 in WT and TRIM24^−/−^ HEK293T cells was assessed by western blotting and qRT-PCR. The mean and SD of the results from three independent experiments are shown (\**P* < 0.05).

**S4 Data.**
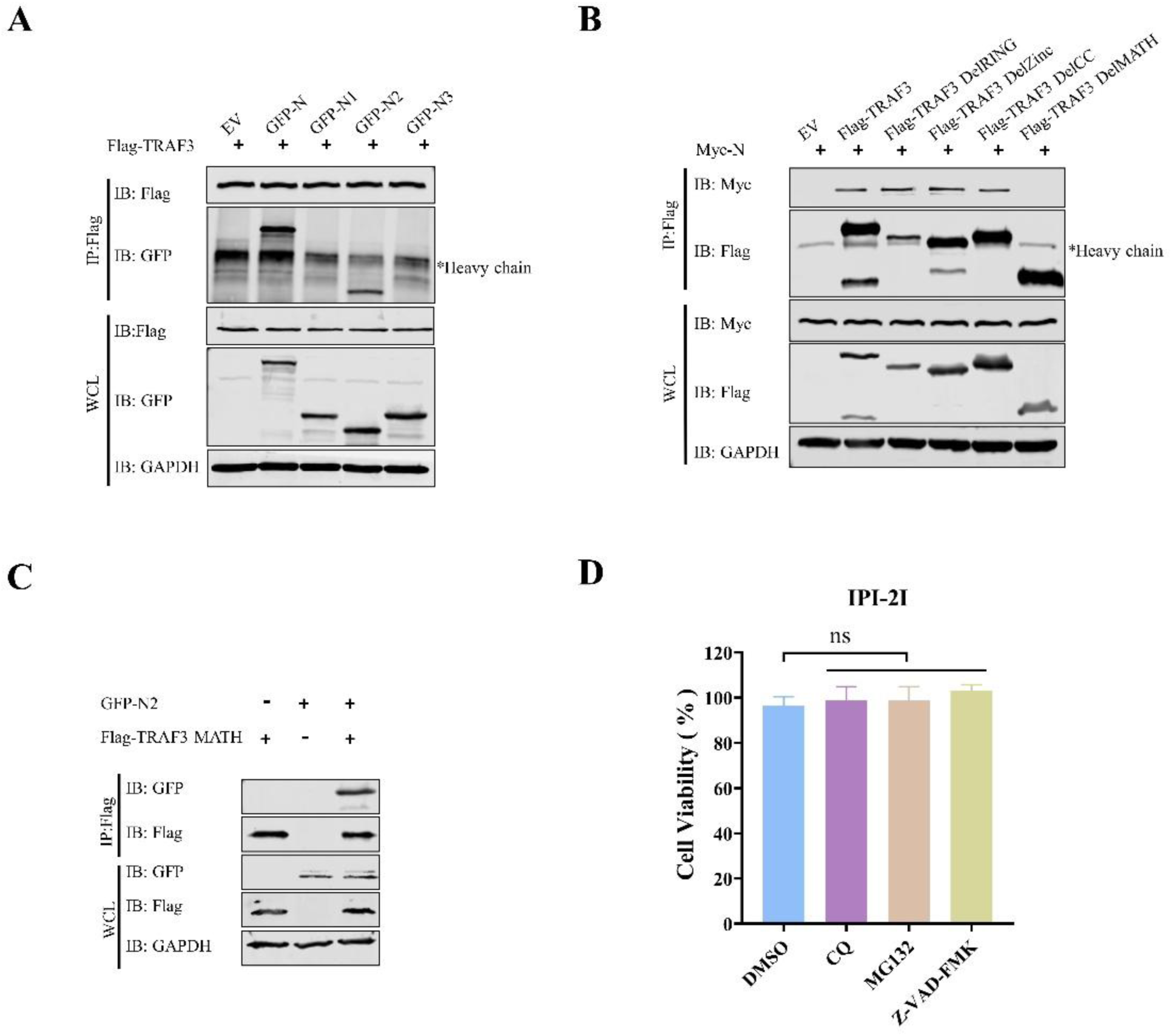
The N2 region of SADS-CoV N protein interacted with the MATH domain of TRAF3. (A) HEK293T cells were co-transfected with plasmids encoding Flag-TRAF3, GFP-N, GFP-N1, GFP-N2, or GFP-N3. Cellular lysates were subjected to immunoprecipitation using Flag Ab. (B) Plasmids encoding TRAF3, DelRING mutant, DelZINC mutant, DelCC mutant, and DelMATH mutant containing Flag tag were co-transfected into HEK293T cells with Myc-N, respectively. Lysates were subjected to immunoprecipitation using Flag Ab. (C) HEK293T cells were co-transfected with plasmids Flag-TRAF3 MATH and GFP-N2. Lysates were subjected to immunoprecipitation using Flag Ab. (D) IPI-2I cells were treated with 20 μM CQ, MG132, or Z-VAD-FMK for 24 h. DMSO-treated cells were used as a control group. Cytotoxicity was detected by CCK-8 assay.

## Acknowledgements

This work was supported by the National Key R&D Program of China (grant no. 2025YFD1800903). The study protocol was approved by the Institutional Animal Care and Use Committee of Harbin Veterinary Research Institute (Approval no. 240624-02-GR) and conducted in accordance with the “Guide for the Care and Use of Laboratory Animals”.

**Author contributions Conceptualization:** Da Shi.

**Data curation:** Miaomiao Zeng, Dakai Liu, Jialin Zhang.

**Funding acquisition:** Da Shi, Li Feng.

**Investigation:** Miaomiao Zeng, Dakai Liu, Jiyu Zhang, Hongyan Shi, Jianfei Chen, Xin Zhang, Zhaoyang Ji, Junyi Su, Xinwei Sun, Xiuwen Li.

**Resources:** Liaoyuan Zhang, Tingshuai Feng.

**Supervision:** Da Shi, Li Feng.

**Writing–original draft:** Miaomiao Zeng, Dakai Liu, Da Shi.

**Writing–review & editing:** Miaomiao Zeng, Dakai Liu, Da Shi.

